# The molecular clock of *Mycobacterium tuberculosis*

**DOI:** 10.1101/532390

**Authors:** F. Menardo, S. Duchêne, D. Brites, S. Gagneux

## Abstract

The molecular clock and its phylogenetic applications to genomic data have changed how we study and understand one of the major human pathogens, *Mycobacterium tuberculosis* (MTB), the causal agent of tuberculosis. Genome sequences of MTB strains sampled at different times are increasingly used to infer when a particular outbreak begun, when a drug resistant clone appeared and expanded, or when a strain was introduced into a specific region. Despite the growing importance of the molecular clock in tuberculosis research, there is a lack of consensus as to whether MTB displays a clocklike behavior and about its rate of evolution. Here we performed a systematic study of the molecular clock of MTB on a large genomic data set (6,285 strains), covering different epidemiological settings and most of the known global diversity. We found that sampling times below 15-20 years were often insufficient to calibrate the clock of MTB. For data sets where such calibration was possible we obtained a clock rate between 1×10^−8^ and 5×10^−7^ nucleotide changes per-site-per-year (0.04 - 2.2 SNPs per-genome-per-year), with substantial differences between clades. These estimates were not strongly dependent on the time of the calibration points as they changed only marginally when we used epidemiological isolates (sampled in the last 40 years) or ancient DNA samples (about 1,000 years old) to calibrate the tree. Additionally, the uncertainty and the discrepancies in the results of different methods were sometimes large, highlighting the importance of using different methods, and of considering carefully their assumptions and limitations.

## Introduction

In 1962, Zuckerland and Pauling used the number of amino-acid differences among hemoglobin sequences to infer the divergence time between human and gorilla, in what was the first application of the molecular clock (Zuckerland and Pauling 1962). Although many at the time found it “crazy” (Morgan 1998), soon the molecular clock was incorporated in Kimura’s neutral theory of molecular evolution (Kimura 1968), and found its place in the foundations of evolutionary biology. Thanks to the improvements of sequencing technologies and statistical techniques, it is now possible to use sequences sampled at different times to calibrate the molecular clock and study the temporal dimension of evolutionary processes in so called measurably evolving populations (Drummond et al. 2003). These advancements have been most relevant for ancient DNA (aDNA), and to study the evolutionary dynamics of pathogen populations, including one of the deadliest human pathogens: *Mycobacterium tuberculosis* (WHO 2018).

In 1994, Kapur and colleagues pioneered molecular clock analyses in MTB: they assumed a clock rate derived from other bacteria and used genetic polymorphisms to infer the age of divergence of different MTB strains (Kapur et al. 1994). Since then, phylogenetic analyses with a molecular clock have been used to estimate the timing of the introduction of MTB clades to particular geographic regions, the divergence time of the MTB lineages, and the age of the most recent common ancestor (MRCA) of the MTB complex (Comas et al. 2013, Bos et al. 2014, Merker et al. 2015, Kay et al. 2015, Brynildsrud et al. 2018, Liu et al. 2018, Rutaihwa et al. 2019). Clock models, together with phylodynamic models in a Bayesian setting have been used to characterize tuberculosis epidemics by determining the time at which outbreaks began and ended (Eldholm et al. 2015, Lee et al. 2015, Folkvardsen et al. 2017, Bainomugisa et al. 2018, Kühnert et al. 2018), establishing the time of origin and spread of drug resistant clades (Cohen et al. 2015, Eldholm et al. 2015, Eldholm et al. 2016, Brynildsrud et al. 2018), and correlating population dynamics with historical events (Pepperell et al. 2013, Merker et al. 2015, Eldholm et al. 2016, Liu et al. 2018, Merker et al. 2018). One example of the potential of molecular clock analyses is the study of Eldholm and colleagues (Eldholm et al. 2016), where the collapse of the Soviet Union and of its health system was linked to the increased emergence of drug resistant strains in former Soviet countries, thus providing insights into the evolutionary processes promoting drug resistance.

A key aspect about estimating evolutionary rates and timescales in microbial pathogens is assessing their clocklike structure. All molecular clock analyses require some form of calibration. In many organisms this consists in constraining internal nodes of phylogenetic trees to known divergence times (for example, assuming codivergence with the host, or the fossil record), but in rapidly evolving pathogens and studies involving aDNA, it is also possible to use sampling times for calibrations (Seo et al. 2002). In the latter approach, the ages of tips of the tree, rather than those of internal nodes are constrained to their collection times. Clearly, the sampling time should capture a sufficient number of nucleotides changes to estimate the evolutionary rate, which will depend on the evolutionary rate of the organism and the extent of rate variation among lineages. Some popular methods to assess such clocklike structure are the root-to-tip regression and the date randomization test (DRT).

While many of the studies inferring evolutionary rates for MTB reported support for a molecular clock (Eldholm et al. 2015, Kay et al. 2015, Eldholm et al. 2016, Folkvardsen et al. 2017, Brynildsrud et al. 2018, Kühnert et al. 2018, Merker et al. 2018, Rutaihwa et al. 2019), some found a lack of clocklike structure (Comas et al. 2013, Bainomugisa 2018, Kühnert et al. 2018), and others assumed a molecular clock without testing whether the data had a temporal structure (Pepperell et al. 2013, Cohen et al. 2015, Merker et al. 2015, Lee et al. 2015, Liu et al. 2018). In all studies where the calibration was based on the sampling time (tip-dating), the clock rate estimates spanned roughly an order of magnitude around 10^−7^ nucleotide changes per site per year. This was in contrast with the results of Comas et al. 2013, where the clock was calibrated assuming co-divergence between MTB lineages and human mitochondrial haplotypes (i.e. internal node calibrations), and was estimated to be around 10^−9^ nucleotide changes per site years. Some lineage 2 (L2) data sets (Eldholm et al. 2016) were found to have a faster clock rate compared to lineage 4 (L4) data sets (Pepperell et al. 2013, Eldholm et al 2015, Folkvardsen et al. 2017, Brynildsrud et al. 2018), while others showed lower clock rates, comparable with L4 (Merker et al. 2018, Rutaihwa et al. 2019). Studies based on aDNA produced slightly lower clock rate estimates (Bos et al. 2014, Kay et al. 2015, Sabin et al. unpublished https://www.biorxiv.org/content/10.1101/588277v1) compared to studies based on modern strains, thus suggesting support for the phenomenon of time dependency of clock rates in MTB (Ho et al. 2011). All these results indicate that different MTB lineages and populations might have different clock rates, and that the age of the calibration points could influence the results of the analyses. Comparing the results of different studies has however a main limitation: the observed differences could be due to different rates of molecular evolution among MTB populations, to methodological discrepancies among studies, or a combination of both.

Here, we assembled a large genomic data set including sequences from all major lineages of MTB (6,285 strains in total, belonging to six human adapted lineages, L1-L6, and one lineage predominantly infecting cattle, *M. bovis*). We then applied the same set of methodologies to the whole data set, to individual lineages and sub-lineages, and to selected local outbreaks, thus ensuring the comparability of the results among different clades and epidemiological settings.

With this systematic approach, we addressed the following questions:

1. Is there a molecular clock in MTB and how do we detect it?
2. What is the clock rate of MTB, and what is its variation among lineages, sub-lineages and individual outbreaks?
3. Are clock rate estimates dependent on the age of the calibration points in MTB?

## Results and Discussion

### Is there a molecular clock in MTB?

Finding evidence of temporal structure is the first step when performing molecular clock analyses (Rieux and Balloux 2016). If there is not enough genetic variation between samples collected at different times, these cannot be used to calibrate the molecular clock, i.e. the population is not measurably evolving. To test the temporal structure of MTB data sets we identified 6,285 strains with a good quality genome sequence, and for which the date of isolation was known (Methods, Sup. Table S1).

We used root to tip regression to evaluate the temporal structure of the whole MTB complex and of the individual lineages (L1-L6 and *M. bovis*) (Rambaut et al. 2016). The root to tip regression is a regression of the root-to-tip distances as a function of sampling times of phylogenetic trees with branch lengths in units of nucleotide changes per site, where the slope corresponds to the rate. Under a perfect clock-like behavior, the distance between the root of the phylogenetic tree and the tips is a linear function of the tip’s sampling year: recently sampled strains are further away from the root than older ones, such that the R^2^ is the degree of clocklike behavior (Korber et al. 2000). We obtained very low values of R^2^ for all lineages (maximum 0.1 for *M. bovis*), indicating a lack of strong clock-like behavior (Sup. Fig. S2). Additionally, we found a weak negative slope for L1, L5 and L6, normally interpreted as evidence for a lack of temporal structure, or overdispersion in the lineage-specific clock rates (Rambaut et al. 2016, Sup. Fig. S2, Sup. Table S3). Negative slope of the regression line can be caused by an incorrect placement of the root (Tong et al.2018). To address this potential problem, we repeated these analyses rooting the trees with an outgroup, we found a negative slope for L1 and L6 and a positive slope for L5, although with an extremely low value of R^2^ (< 0.01). These results indicate that the negative slope of L1 and L6 and the low R^2^ values of the three data sets are not due to an incorrect placement of the root (Sup. Fig. S4).

Since root to tip regression can be used only for exploratory analyses and not for formal hypothesis testing (Rambaut et al. 2016), we performed a date randomization test (DRT). The DRT consists in repeatedly reshuffling the date of sampling among taxa and then comparing the clock rate estimates among the observed and reshuffled data sets (Rieux and Balloux 2016). If the estimation obtained from the observed data does not overlap with the estimations obtained from the randomized data sets, we can conclude that the observed data has a stronger temporal signal than expected by chance, such that there is statistically significant clocklike structure (Rieux and Balloux 2016). Usually the DRT is implemented in a Bayesian phylogenetic setting, however, considering the size and the number of data sets included in this study, an excessive amount of computation would be required. To overcome this problem, we estimated the clock rate with the least-squared dating method implemented in LSD (To et al. 2015). The advantage of this method is that it is orders of magnitude faster than fully Bayesian approaches, and can therefore be used on data sets with thousands of taxa and with more randomizations compared to the10-20 typically used in a Bayesian setting (Duchene et al. 2018). A limitation of least squares dating is that it typically assumes a single tree topology and vector of branch lengths, and a strict clock (i.e. all branches have the same clock rate). However, a simulation study showed that maximum likelihood trees produced similar estimates compared to the true topology, and that it is robust to uncorrelated variation of the clock rate among branches in the phylogeny (To et al. 2015, Duchene et al. 2016 a, Duchene et al. 2018).

For each data set, we reshuffled the year of sampling among tips 100 times and estimated the clock rate of observed and randomized data sets with LSD. All eight data sets except L5 and L6 passed the DRT (Methods, Sup. Fig. S2, Sup. Table S3). L5 and L6 are the two lineages with the lowest sample size, 117 and 33 strains, respectively. Moreover most strains were sampled in a short temporal period compared to the other lineages (Sup. Figs. S5-S10). It is likely that with additional strains sampled across a larger time period, L5 and L6 will also show evidence for a molecular clock.

We complemented the analysis described above with a Bayesian phylogenetic analysis in Beast2 (Bouckaert et al. 2014). Since this is computationally expensive, we reduced the large data sets (MTBC, L1, L2, L4 and *M. bovis*) to 300 randomly selected strains. For each data set we selected the best fitting nucleotide substitution model identified with jModelTest 2 (Darriba et al. 2012). For this first analysis, we assumed a coalescent constant population size prior, used a relaxed clock model, and a 1/x prior for the clock rate, constrained between 10^−10^ and 10^−5^ nucleotide changes per site per year. This interval spans the range of clock rates proposed for *M. tuberculosis* and for most other bacteria (Duchene et al 2016 b, Eldholm et al. 2016). We observed that for all data sets the posterior was much more precise (with a narrow distribution) than the prior, thus indicating that the data was informative (Drummond et al. 2006). Again, the only exceptions were L5 and L6, where the posterior distribution was flat, ranging between 10^−10^ and 10^−7^ nucleotide changes per site per year, confirming the lack temporal structure of these two data sets (Sup. Fig. S2).

We repeated these analyses on 23 sub-lineages and 7 outbreaks and local populations to test whether we could detect a temporal structure also in smaller, less diverse data sets. With this sub-sampling scheme, we could compare the results among different clades, among outbreaks with different epidemiological characteristics, and among local outbreaks and global data-sets (see Methods). We found that 11 sub-lineages and 5 local populations passed the DRT (Sup. Table S3, Sup. Figs. S5-S8 and S11-S13).

All the data sets that failed the DRT had less than 350 genomes, or were composed of strains sampled in a temporal range of 20 years or less. Additionally, only two of the ten data sets sampled across less than 15 years, and three of the twelve data sets with less than 100 strains passed the DRT (Fig. 1; Sup. Table S2), indicating that large sample sizes and wide temporal sampling windows appear to be necessary to obtain reliable estimates of evolutionary rates and timescales in MTB. Conversely, the number of polymorphic positions and the genetic diversity measured with Watterson’s estimator did not correlate with the outcome of the DRT (Sup. Fig S14).

**Figure 1.**
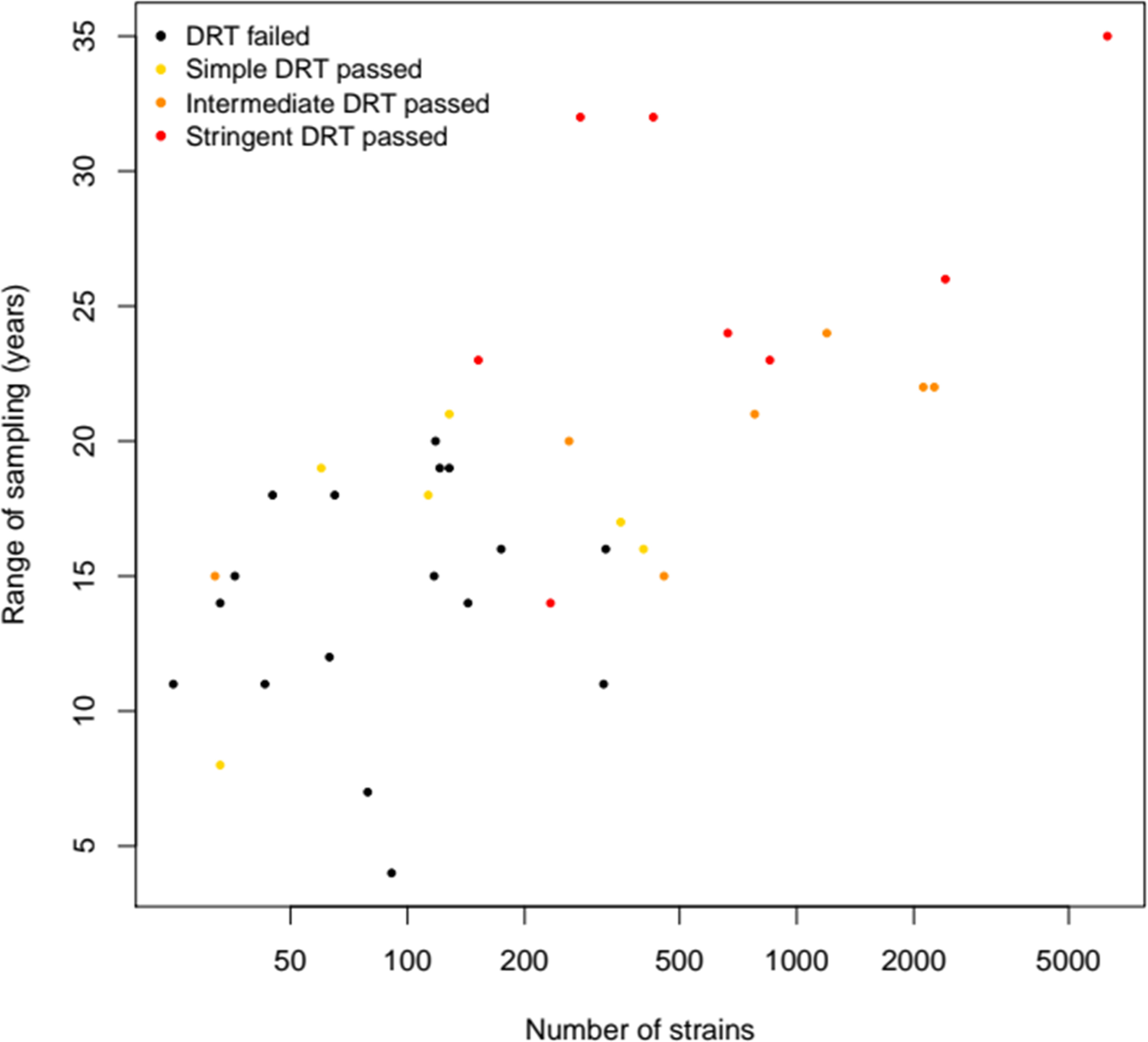
Results of the DRT for all data sets ordered by size and temporal range. Data sets with fewer strains sampled in a shorter period of time tended to fail the DRT.

Among the three methods generally used to study the temporal structure of a data set, the root to tip regression resulted in a negative slope, and therefore failed to detect the temporal structure of some of the data sets that passed the DRT (i.e. L1, L4.1.2 and L1.1.1). Nevertheless, root to tip regression can be useful to identify data sets where the temporal signal comes from a single strain, or a few strains (see below). Comparing prior and posterior distributions of the clock rates was also useful to detect the presence of temporal structure, although this was not always in agreement with the results of the DRT: some of the data sets that did not pass the DRT (e.g. L2.2.1_nc2, Trewby 2016) had a posterior distribution of the clock rate more distinct from the prior than some of the data set that passed the DRT (e.g. L1.1.1, L1.2.1 and L1.2.2) (Sup. Figs. S5 and S7-S8, Sup. Table S3). A possible reason for this could be that LSD and Beast have different statistical power with different data sets. Additionally, in some cases the deviation of the posterior distribution of the clock rate from the prior could be an artifact caused by tree prior misspecification, and not the result of genuine temporal structure (Möller et al. 2018).

### Sensitivity of the clock rate estimates to the model assumptions

In Bayesian analyses, different models and priors are based on different assumptions about the evolutionary processes, and can thus influence the results (Bromham et al. 2018). Often different sets of assumptions are tested in a Bayesian framework by comparing their marginal posterior probability with the Bayes factor, and the most likely model is then chosen to estimate the parameters of interest (Bromham et al. 2018). Given the size and number of the data sets considered in this study, it is not possible to assess the relative fit of many competing models for all data sets. However, model misspecification can result in biased estimates. It is therefore important to investigate the robustness of the results to different models and priors.

We repeated the Bayesian analysis using a uniform prior instead of the 1/x prior on the clock rate. We ran a Beast analysis sampling from the priors and found that the uniform prior was biased towards high clock rates and put most weight on rates between 10^−6^ and 10^−5^ nucleotide changes per site per year (Sup. Fig. S15). For all data sets, we compared the posterior distribution of the clock rate obtained with the two different priors (Sup. Figs. S16-S18, Sup. Table S3).

Some data sets showed hardly any difference (e.g. MTBC, L1, L2, L3, L4 etc.), indicating that the data was informative and that the data set had a strong temporal structure. However, this did not always correlate with the results of the DRT. For example, the subset of 300 strains of L2 and the data set Trewby 2016 did not pass the DRT but showed a distinct posterior distribution that was not sensitive to the prior choice. Other data sets, including three that passed the DRT by a small margin (L1.1.1, L1.2.1 and L1.2.2), were more sensitive to the prior choice and resulted in two distinct posterior distributions, indicating a weaker temporal structure (Sup. Fig. S8).

An additional assumption of the phylogenetic model that can influence the results of molecular clock analyses is the tree prior (also known as demographic model). We tested the sensitivity to the tree prior by repeating the analysis with an exponential population growth (or shrinkage) prior instead of the constant population size. For this analysis, we used the 1/x prior on the clock rate and we considered only the data sets that passed the DRT (21 data sets). The constant population model is a specific case of the exponential growth model (when the growth rate is equal to zero). Therefore, if the 95% Highest Posterior Density interval (HPD) of the growth rate does not include zero, we can conclude that the data reject a demographic model with constant population size. We found that 14 data sets rejected the constant population size model, and that all of them had positive growth rates (Sup. Table S3). The three data sets that were found to be sensitive to the prior on the clock rate were also sensitive to the tree prior, confirming their low temporal structure and information content, while the results for all other data sets were only moderately influenced by the tree prior (Sup. Figs. 19-20, Sup. Table S3).

Overall, we found that, except for three data sets (L1.1.1, L1.2.1 and L1.2.2), the clock rate estimates were robust to different priors of the clock rate and to different demographic models. To compare the clock rates of different data sets, we report the analysis with the 1/x prior on the clock rate because the uniform prior can bias the estimates upward. For data sets that showed evidence against the constant population size model (95% HPD of the growth rate not including zero), we report the results of the analysis with the exponential population growth, and for the others, we report the results of the analysis with constant population size.

### What is the clock rate of MTB, and what is its variation among lineages, sub-lineages and outbreaks?

We found that the point estimates of all data sets where we detected temporal structure range between 2.86×10^−8^ (L3 Beast) and 4.82×10^−7^ (Eldholm 2016 Beast) nucleotide changes per site per year. While some data sets had a low range of the 95% confidence interval (CI), reaching the hard limit imposed by LSD of 10^−10^, most of the CI and 95% highest posterior density intervals (HPD) are included between 10^−8^ and 5×10^−7^ (Fig. 2 and Sup. Table S3). This range encompasses previous estimates obtained with epidemiological samples and aDNA and is among the lowest in bacteria, thus supporting our conclusion from above: tip-dating with MTB requires samples collected over a long period of time because of the slow clock rate.

**Figure 2.**
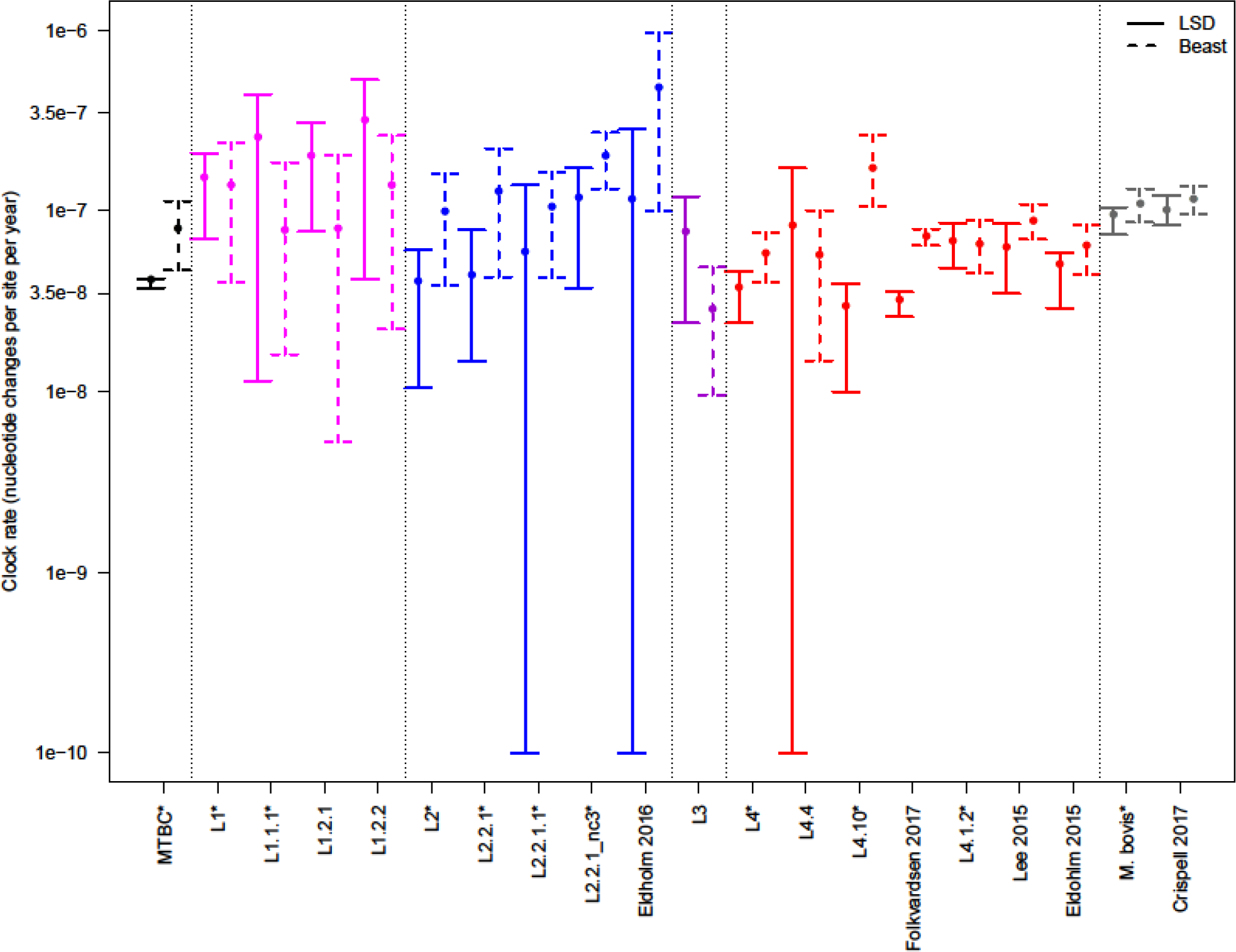
Estimated clock rates of all lineages, sub-lineages and local data sets that passed the DRT. Solid lines represent the 95% confidence interval estimated with LSD, dashed lines represent the 95% highest posterior density (HPD) estimated by Beast (the larger dot is the median of the posterior distribution). We show here the results of the Beast analysis with the 1/x prior on the clock rate. For data sets that rejected the constant population size, we show the result obtained with the exponential population growth prior, for the other data sets we show the results obtained with the constant population size prior. Data sets marked with * have been reduced to 300 randomly picked strains for the Beast analysis.

There was one notable exceptions to the pattern described above: the data sets L4_nc which showed a much higher clock rate estimate compared to all other data sets included in this study (~10^−6^; Sup. Table S3). However, this is most likely an artifact: 1) L4_nc is the smallest among all considered data sets, with 32 strains. 2) Most strains are identical or nearly so, collected in the same year, and form a monophyletic clade (Sup. Figs. S9 and S21). It is known that data sets with a high degree of temporal and phylogenetic clustering can pass the DRT also when they do not have temporal structure (Duchene et al. 2015). 3) The root to tip regression suggests that the temporal signal comes from one single strain in L4_nc (Sup. Fig. S7). We therefore excluded the L4_nc data set from further analyses.

Our results suggest that different lineages of MTB have different clock rates, for example most L1 data sets had point estimates higher than most L4 data sets, although the CI and HPD were often overlapping. The point estimates indicate that the clock rate of L1 is more than double the clock rate of L4: two average L1 strains are expected to differ by 12 SNPs after ten years of divergence, while two average L4 strains will differ by 5 SNPs after the same period of time. This was supported by the results of both LSD, where the 95% CI of L1 and L4 did not overlap, and Beast, where the 95% HPD overlapped partially, but the two posterior distributions showed distinct peaks (Fig. 2, Sup Table S3, Sup. Fig. S22). A practical implication of these results pertains to the widespread use of SNP distances to identify ongoing transmission in MTB epidemiological studies. Usually, recent transmission is postulated when two or more strains differ by a number of SNPs below a certain threshold (Hatherell et al. 2016). However this approach will result in systematically lower levels of transmission for clades with faster rates of molecular evolution. For example, a recent study reported low transmission rates of L1 compared to L2 and L4 in Vietnam (Holt et al. 2018), which could partially be explained by a faster clock rate of L1, opposed to reduced ongoing transmission.

When considering the results of Beast, also L2 had a higher clock rate compared to L4, and all data sets included in the sub-lineage L2.2.1 showed a faster clock rate compared to the complete L2 data set (Fig. 2). The sub-lineage L2.2.1 includes the so called “modern Beijing” family, which was shown to be epidemiologically associated with increased transmission, virulence and drug resistance (Glynn et al. 2006, Hanekom et al. 2010, de Steenwinkel et al. 2012, Ribeiro et al. 2014, Holt et al. 2018, Wiens et al. 2018), and to have a higher mutation rate compared to L4 strains (Ford et al. 2013). However, the LSD estimates for L2.2.1 and for its sub-lineages, despite showing the same trend of Beast, support a lower clock rate compared to Beast, and have large confidence intervals, overlapping with the results of L2 and L4 (Fig. 2).

Further evidence of among-lineage variation is provided by the results of the Bayesian analyses, where for most data sets we obtained coefficients of variation (COV) with a median of 0.2 – 0.3, and not abutting zero (Sup. Table S3), thus rejecting the strict clock (Drummond et al. 2006).

Taken together, these results indicate that there is a moderate variability among the current rate of molecular evolution of different MTB lineages, which could be caused by different mutation rates as it was reported for L2 and L4 (Ford et al. 2013), and support the idea that the inference of transmission in MTB should move from the use of SNP distances to methods that incorporate information about the molecular clock (Stimson et al. 2019).

In our analysis we included two outbreaks caused by strains belonging to the same sub-lineage (L4.1.2; Eldholm et al. 2015, Lee et al. 2015). This gives us the opportunity to compare the molecular clock of clades with a similar genetic background in different epidemiological settings. The Eldholm 2015 data set is a sample of an outbreak in Argentina, in which resistance to multiple antibiotics evolved several times independently (Eldholm et al. 2015). The Lee 2015 data set represents an outbreak of drug susceptible strains in Inuit villages in Québec (Canada). The clock rates of these two data sets were highly similar (95% CI and HPD ranging between 5.07×10^−8^ and 8.88×10^−8^ for all analyses; Fig. 2, Sup. Table S3) thus suggesting that, at least in this case, different epidemiological characteristics, including the evolution of antibiotic resistance, do not have a large impact on the rate of molecular evolution of MTB.

### A faster clock for the ancestor of *M. bovis*?

We showed that current clock rates are moderately different among different data sets. A different question is whether the clock rate was constant during the evolutionary history of the MTB complex. When looking at the phylogenetic tree of the MTB complex, rooted with the genome sequence of *M. canettii*, one notices that strains belonging to different lineages, despite being all sampled in the last 40 years, have different distances from the root (Fig. 3). For example, since their divergence from the MRCA of the MTB complex, the two *M. africanum* lineages (L5 and L6) and especially *M. bovis*, accumulated more nucleotide changes than the lineages belonging to MTB sensu stricto (L1-L4; Fig. 3). Additionally, all methods (root to tip regression, LSD and Beast) if used without an outgroup, placed the root on the branch between *M. bovis* and all other lineages, while rooting the tree with the outgroup *M. canettii* placed the root on the branch connecting MTB *sensu stricto* with *M. africanum* (L5 and L6) and *M.bovis.* The different root placement affects the clock rate estimation only moderately (4.16×10^−8^ LSD analysis without outgroup, 5.59×10^−8^ LSD analysis with outgroup, Sup. Table S3), but it is a further indication of the variation of the rate of molecular evolution during the evolutionary history of the MTB complex. The observation that all *M. bovis* strains, despite having a clock rate similar to all other data sets, have a larger distance from the root of the MTB complex tree compared to other lineages is intriguing, and could be explained by a faster rate of molecular evolution of the ancestors of *M. bovis* (Figs. 2-3). It is believed that *M. bovis* switched host (from human to cattle) (Brosh et a. 2002, Mostowy et al. 2002, Brites et al. 2018), and it is possible that during the adaptation to the new host several genes were under positive selection, thus leading to an increase in the accumulation of substitutions in the *M. bovis* genome. Another possibility is that the ancestor of *M. bovis* experienced a period of reduced population size, a bottleneck, and as a consequence, slightly deleterious mutations were fixed by genetic drift, resulting in a faster clock rate compared to larger populations where selection is more efficient in purging deleterious mutations (Ohta 1987, Bromham and Penny 2003).

**Figure 3.**
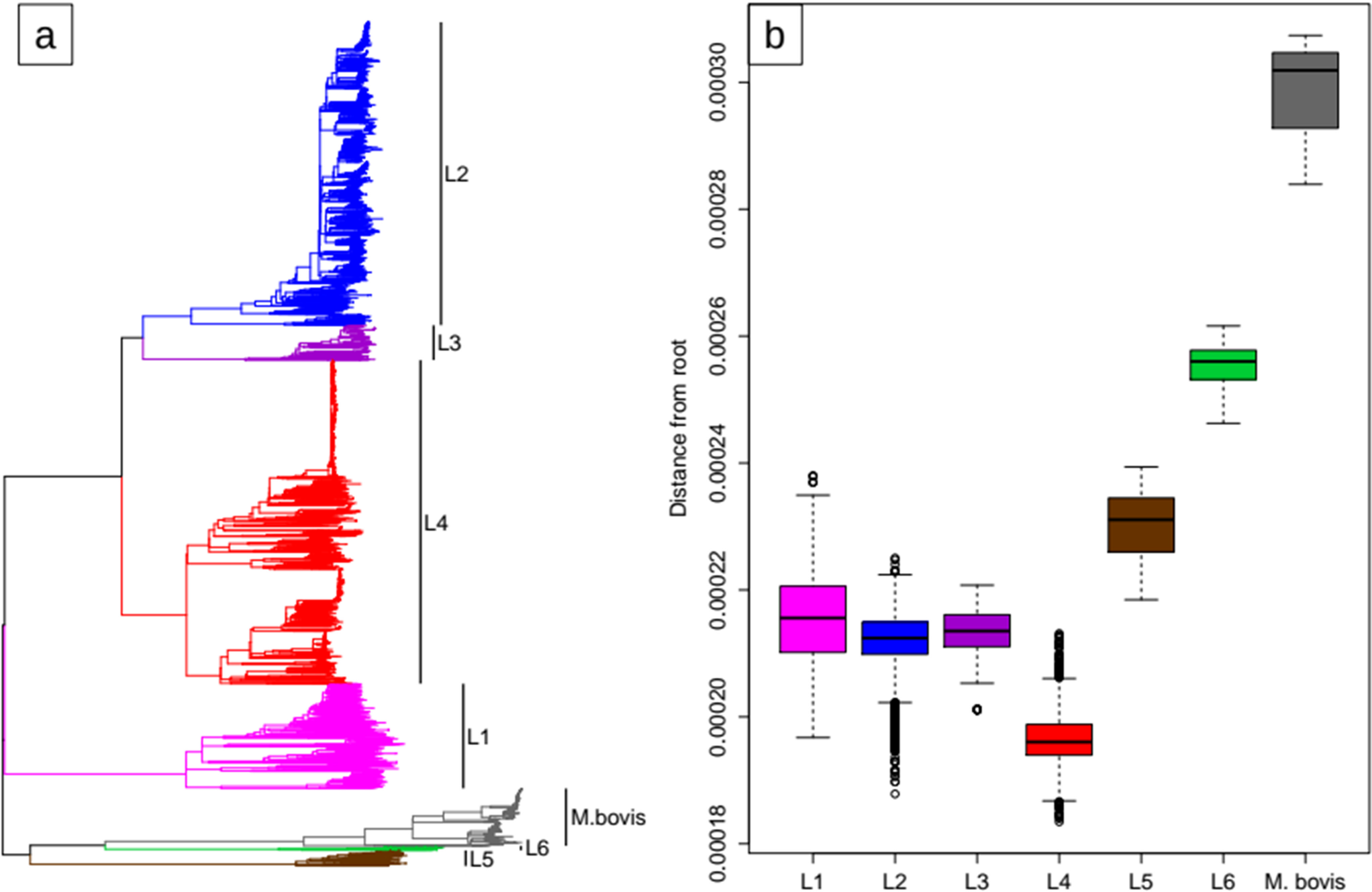
**a)** The Maximum Likelihood tree of the 6,285 strains considered in this study rooted with the genome sequence of *M. canetti*. **b)** Phylogenetic distance from the root (expected nucleotide changes per site) of MTB strains by lineage

### Time dependency of the clock rate

It has been suggested that in MTB, as in other organisms, the clock rate estimation is dependent on the age of the calibration points (Ho et al. 2005, Ho et al. 2011, Comas et al. 2013, Duchene et al. 2014, Duchene et al. 2016 b), and that using recent population-based samples could result in an overestimation of the clock rate, because these samples include deleterious mutations that have not yet been purged by purifying selection. However, the validity of the time dependency hypothesis has been contested in general (Emerson and Hickerson 2015), and for MTB in particular (Pepperell et al. 2013). Here we used an approach similar to Rieux et al. (2014) and tested whether the time dependency hypothesis was supported by our data. We repeated the analyses presented above, only this time we included the aDNA genome sequences of three MTB strains obtained from Precolumbian human remains from Peru (Bos et al. 2014). If the clock rate estimates depend on the age of the calibration points, adding ancient genomes should result in lower clock rates. We performed this analysis with LSD, using the complete data set (6,285 strains), and with Beast, using the sub-sample of 300 randomly selected strains described above, and an additional independent random sub-sample of 500 strains (Methods).

With LSD, adding the aDNA samples resulted in a slightly faster clock rate, conversely all the analyses performed with Beast resulted in marginally slower clock rates when the aDNA samples were included (Table 1). These results indicate that the effect of the age of the calibration points on the clock rate is modest, and they are corroborated by the observation that MTB mutation rates in vitro and in vivo, estimated with fluctuation assays and resequencing of strains infecting macaques, are remarkably similar to the clock rates obtained in our study (~ 3×10^−8^ - 4×10^−7^; Ford et al. 2011).

**Table1.** Results of LSD and Beast for the MTB complex with and without aDNA samples. With Beast we performed three different analyses, two using a sub-sample of 300 strains and different priors on the clock rate (1/x and uniform), and one using a sub-sample of 500 strains (Methods, Sup. Table S3))

| Dataset | Clock rate |
| --- | --- |
| MTBC 6,285 + M. canettii (LSD) | 5.59E-08 (4.12E-08, 6.17E-08) |
| MTBC 6,285 + M. canettii + aDNA<br>(LSD) | 6.93E-08 (5.48E-08, 8.42E-08) |
| MTBC 300 (Beast 1/x) | 8.2254E-08 (4.964E-08, 1.141E-07) |
| MTBC 300 +aDNA (Beast 1/x) | 7.3978E-08 (4.648E-08, 1.019E-07) |
| MTBC 300 (Beast unif) | 8.26E-08 (4.82E-08, 1.14E-07) |
| MTBC 300 +aDNA (Beast unif) | 7.1794E-08 (4.157E-08, 9.851E-08) |
| MTBC 500 (Beast unif) | 6.08E-08 (4.21E-08, 8.07E-08) |
| MTBC 500 +aDNA (Beast unif) | 5.20E-08 (3.41E-08, 7.12E-08) |

The aDNA samples considered in this study are not optimal to test the time dependency hypothesis because they belong to the *M. pinnipedii* clade of the MTB complex (Bos et al. 2014). The modern strains of this lineage are rarely sampled, because they are infecting seals and sea lions rather than humans. The only additional aDNA samples available for MTB are L4 samples isolated from 18^th^ century Hungarian mummies (Kay et al. 2013), however these samples are a mix of strains with different genotypes, and cannot be easily integrated with the data and pipelines used in this study. Additional aDNA samples from older periods and belonging to other lineages are necessary to better investigate the time dependency hypothesis in MTB. Recently, Sabin and colleagues (Sabin et al. unpublished: https://www.biorxiv.org/content/10.1101/588277v1) reported the sequencing of a high quality MTB genome from the 17^th^ century, this data will contribute to the investigation of the time dependency hypothesis in MTB.

### Dating MTB phylogenies

In most cases, the goal of molecular clock studies is not to estimate the clock rates, but rather the age of the phylogenetic tree and of its nodes. Conceptually, this means extrapolating the age of past events from the temporal information contained in the sample set. If we exclude the few aDNA samples that are available (Bos et al. 2014, Kay et al. 2015), all MTB data sets have been sampled in the last 40 years. It is therefore evident that the age estimates of recent shallow nodes will be more accurate than medium and deep nodes. In part, this is reflected in the larger CI and HPD of the age of ancient nodes compared to more recent ones. Extrapolating the age of trees that are thousands of years old with contemporary samples is particularly challenging, because the observed data captures only a small fraction of the sample’s evolutionary history, and these are the cases where aDNA samples are most valuable.

Nevertheless, the age of the MRCA of the MTB complex and of its lineages is highly relevant to understand the emergence and evolution of this pathogen and a debated topic (Wirth et al. 2008, Comas et al. 2013, Bos et al. 2014). The LSD analyses on the tree rooted with *M. canettii* estimated the MRCA of the MTB complex to be between 2,828 and 5,758 years old (Sup. Table S3). These results are highly similar to the ones of Bos and colleagues (2,951 – 5,339) which were obtained with Bayesian phylogenetics and a much smaller sample size (Bos et al. 2014). These estimates should be taken with caution because of the intrinsic uncertainty in estimating the age of a tree that is several thousands of years old, calibrating the molecular clock with the sampling time of modern strains and only 3 aDNA samples. A more approachable question is the age of the MRCA of the individual MTB lineages. Here we can consider the results of four different analyses: the LSD and Beast analyses on the individual lineages (L1-L4, and *M. bovis*), and the LSD and Beast analyses on the complete MTB complex (including the aDNA samples), from which the age of the MRCA of the lineages can be extracted (L1-L6, and *M. bovis*). When we combined all these results, merging the CI and HPD, we obtained an estimate of the age of the MTB lineages which accounts for the uncertainty intrinsic in each analysis, but also for the differences among inference methods and models, thus providing a more conservative hypothesis. In all our analyses, the point estimates of the age of all lineages resulted to be at most 2,500 years old, and the combined CI and HPD extended to a maximum of 11,000 years ago for L2 (95% CI of the LSD analysis; Fig. 4, Sup. Table S23). The large CI of L2 was maybe due to among-lineage variation of the clock rate in L2, as discussed above. While L5, L6 and *M. bovis* have younger MRCAs and narrower confidence intervals, we should note that for these lineages the sampling is much less complete compared to L1-L4, and it is possible that further sampling will add more basal strains to the tree, thus resulting in older MRCAs. For the other lineages, where the sampling is more representative of the global diversity, the confidence intervals of the age of the MRCAs extend over several thousands of years, and the point estimates of the four analyses spread over 1,000 – 2,000 years. This shows that we should be very careful when interpreting the results of tip dating in MTB, especially if our goal is to estimate the age of ancient nodes such as the MRCAs of MTB lineages. Conservative researchers might want to use different methods; several model and prior combinations should be formally tested in Beast, and the final results can be combined in one range providing an estimation of the uncertainty of the clock rate and of the age of some specific node of the tree.

**Figure 4.**
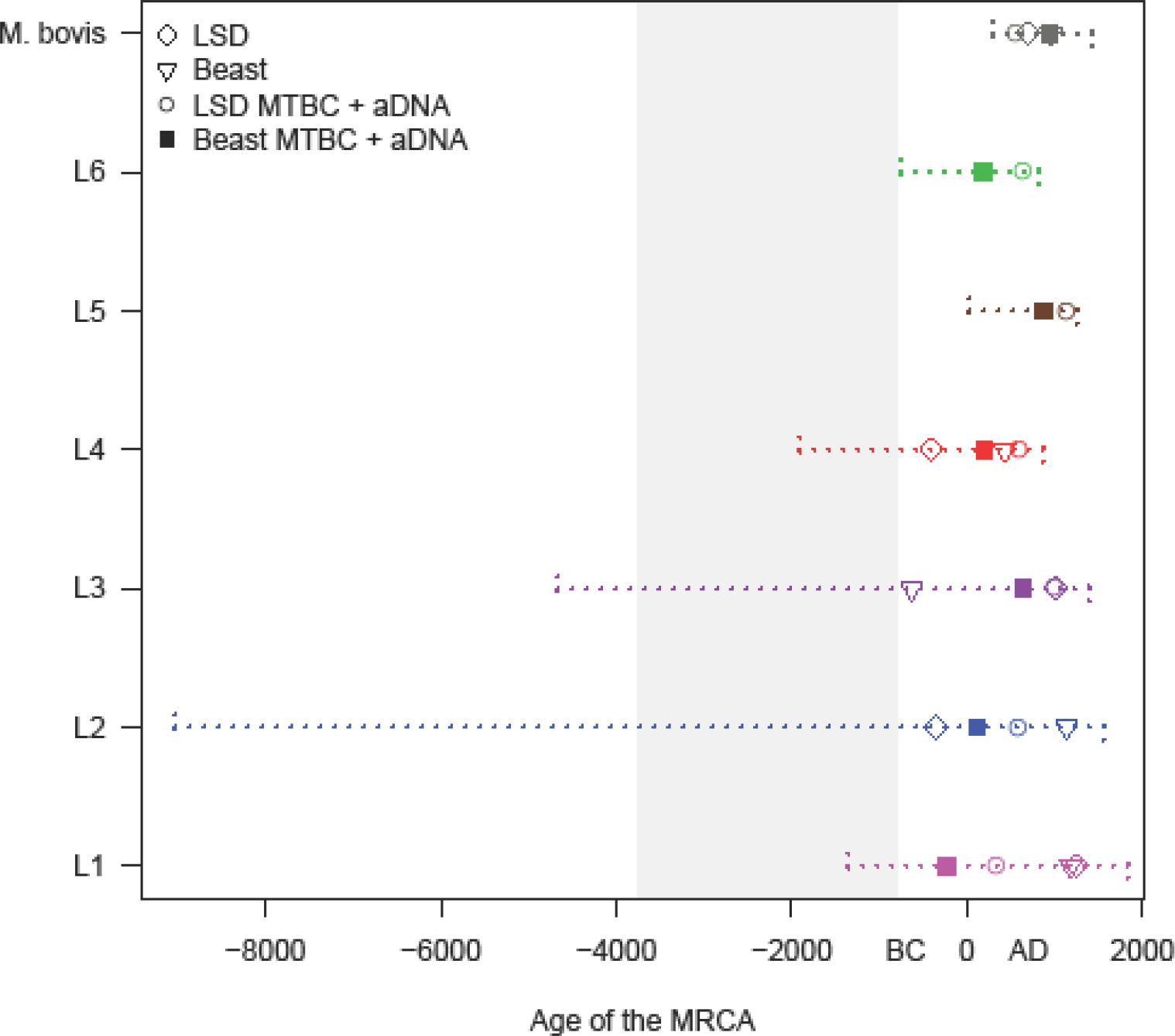
The inferred age of the MTB lineages. LSD : results of the LSD analysis performed on the individual lineages. Beast: results of the Beast analysis performed on the individual lineages (median values). LSD MTBC + aDNA: results of the LSD analysis performed on the complete data set of 6.285 strains + 3 aDNA samples, the age of the MRCA of the individual lineages was identified on the calibrated tree. Beast MTBC + aDNA: results of the Beast analysis performed on the random subsample of 500 strains + 3 aDNA samples, the age of the MRCA of the individual lineages was identified on the calibrated tree (median values). The confidence intervals were obtained merging the 95% CI and HPD of all analyses. The shaded area represents the age of the MRCA of the MTB complex obtained with the LSD analyses (with and without aDNA, the two 95% CI were merged). For L5 and L6 we report only the age inferred on the complete MTB complex tree, because when analyzed individually these two data sets showed a lack of temporal structure (they failed the DRT).

Altogether our results highlight the uncertainty of calibrating MTB trees with tip-dating, they nevertheless support the results of Bos et al. 2014 that found the MRCA of the MTB complex to be relatively recent, and not compatible with the out of-Africa-hypothesis (Wirth et al. 2008, Comas et al. 2013) in which the MTB lineage differentiated in concomitance with the dispersal of *Homo sapiens* out of Africa, about 70,000 years ago. Dating analyses based on DNA samples can only reconstruct the evolutionary history of the data set as far back as the MRCA of the sample. It is possible that in the future new lineages will be sampled, and the MTB phylogeny will be updated moving the MRCA further in the past. Additionally, it is also possible that extinct lineages were circulating and causing diseases much earlier that the MRCA of the strains that are circulating now. This hypothesis is supported by the detection of molecular markers specific for MTB in archeological samples (reviewed in Brites and Gagneux 2015), the oldest of them in a bison’s bone about 17,500 years old (Rothschild et al. 2001). Several such studies directly challenge the results of tip-dating presented here because they reported molecular markers specific to MTB lineages in archeological samples that predate the appearance of those lineages as estimated by tip dating (Taylor et al. 2007, Hershkovitz et al. 2008, Nicklisch et al. 2012). However, there is a controversy regarding the specificity of some of the used markers, and the potential contamination of some of the samples by environmental mycobacteria (Wilbur et al. 2009, Donoghue et al. 2009).

Whole genome sequences from additional aDNA samples are needed to reconcile these two diverging lines of evidence. Ideally they should belong to different lineages, span different periods, and include samples older than the currently available aDNA from Peruvian human remains.

## Methods

### Bioinformatic pipeline

We identified 21,734 MTB genome sequences from the sequence read archive (Sup. Table S24). All genome sequences were processed similarly to what was described in Menardo et al. (2018). We removed Illumina adaptors and trimmed low quality reads with Trimmomatic v 0.33 (SLIDINGWINDOW:5:20) (Bolger et al. 2014). We excluded all reads shorter than 20 bp and merged overlapping paired-end reads with SeqPrep (overlap size = 15) (https://github.com/jstjohn/SeqPrep). We mapped the resulting reads to the reconstructed ancestral sequence of the MTBC (Comas et al. 2013) using the mem algorithm implemented in BWA v 0.7.13 (Li and Durbin 2009). Duplicated reads were marked by the MarkDuplicates module of Picard v 2.9.1 (https://github.com/broadinstitute/picard). We performed local realignment around Indel with the RealignerTargetCreator and IndelRealigner modules of GATK v 3.4.0 (McKenna et al. 2010). We used Pysam v 0.9.0 (https://github.com/pysam-developers/pysam) to exclude reads with alignment score lower than (0.93*read_length)-(read_length*4*0.07)): this corresponds to more than 7 miss-matches per 100 bp. We called SNPs with Samtools v 1.2 mpileup (Li 2008) and VarScan v 2.4.1 (Koboldt et al. 2012) using the following thresholds: minimum mapping quality of 20; minimum base quality at a position of 20; minimum read depth at a position of 7X; minimum percentage of reads supporting the call 90%; no more than 90%, or less than 10% of reads supporting a call in the same orientation (strand bias filter). SNPs in previously defined repetitive regions were excluded (Comas et al. 2013). We excluded all strains with average coverage < 15 X. Additionally, we excluded genomes with more than 50% of the SNPs excluded due to the strand bias filter, and genomes with more than 50% of SNPs with a percentage of reads supporting the call included between 10% and 90%. We filtered out genomes with phylogenetic SNPs belonging to different lineages or sub-lineages (only for L4) of MTB, as this is an indication that a mix of strains could have been sequenced. To do this, we used the diagnostic SNPs obtained from Steiner et al. 2014 and Stucki et al. 2016 for L4 sub-lineages. We excluded all strains for which we could not find the date of isolation 1) in the SRA meta-information, 2) in the associated publications, 3) from the authors of the original study after inquiry. We divided all remaining strains by lineage (L1-L6 and *M. bovis*), and excluded strains with a number of called SNPs deviating more than three standard deviations from the mean of the respective lineage. We built SNPs alignments for all lineages including only variable positions with less than 10% of missing data. Finally, we excluded all genomes with more than 10% of missing data in the alignment of the respective lineage. After all filtering steps, we were able to retrieve 6,285 strains with high quality genome sequences and an associated date of sampling (Sup. Table S1).

### Dataset subdivision

To perform a systematic analysis of the molecular clock in MTB we considered different data sets:

1. the complete data set (6,285 strains)
2. the different lineages of MTB (L1, L2, L3, L4, L5, L6, *M. bovis*)
3. the sub-lineages of L1 (L1.1.1, L1.1.1.1, L1.1.2, L1.1.3, L1.2.1 and L1.2.2) and L2 (L2.1, L2.2.1, L2.2.2 and L2.2.1.1) as defined by Coll et al. 2014; the sub-lineages of L4 (L4.1.1, L4.1.2, L4.1.3, L4.4, L4.5, L4.6.1 and L4.10) as defined by Stucki et al. 2016. Additionally, we identified two L4 clades that were not classified by the diagnostic SNPs of Stucki et al. 2016 (L4_nc and L4.1_nc, respectively, included into L4.6.2 and L4.1.2 as defined by Coll et al. 2014), and three sub-clades of L2.2.1 that were not previously designated as sub-lineages (L2.2.1_nc1, L2.2.1_nc2 and L2.2.1_nc3) (Supplementary Figs. S11-S13).
4. Selected data sets representing outbreaks or local populations that have been used for molecular clock analyses in other studies

Lee et al. 2015 - Mj clade outbreak among a Inuit population in Canada (L4)
Eldholm et al. 2015 - Multi-drug resistant outbreak in Argentina (L4)
Eldholm et al. 2016 – Afghan family outbreak in Oslo (L2)
Trewby et al. 2016 – *M. bovis* in Northern Ireland
Crispell et al. 2017 – *M. bovis* in New Zealand
Folkvardsen et al. 2017 - C2/1112-15 outbreak in Denmark (L4)
Bainomugisa et al. 2018 – Multi-drug resistant outbreak on Daru island in PNG (L2)

### LSD analysis

For all data sets, we assembled SNPs alignments including variable positions with less than 10% of missing data. We inferred phylogenetic trees with raxml 8.2.11 using a GTR model (-m GTRCAT -V options). Since the alignments contained only variable positions, we rescaled the branch lengths of the trees rescaled_branch_length = ((branch_length * alignment_lengths) / (alignment_length + invariant_sites)), Duchene and colleagues (Duchene et al. 2018) showed that this method produced similar results compared to ascertainment bias correction. We then used the R package ape (Paradis et al. 2018) to perform root to tip regression after rooting the trees in the position that minimizes the sum of the squared residuals from the regression line. Root to tip regression is only recommended for exploratory analyses of the temporal structure of a dataset and it should not be used for hypothesis testing (Rambaut et al. 2016). A more rigorous approach is the date randomization test (DRT)(Ramsden et al. 2008), in which the sampling dates are reshuffled randomly among the taxa and the estimated molecular clock rates estimated from the observed data is compared with the estimates obtained with the reshuffled data sets. This test can show that the observed data has more temporal information that data with random sampling times. For each dataset, we used the least square method implemented in LSD v0.3-beta (To et al. 2015) to estimate the molecular clock in the observed data and in 100 randomized replicates. To do this, we used the QPD algorithm allowing it to estimate the position of the root (option -r a) and calculating the confidence interval (options -f 100 and -s). We defined three different significance levels for the DRT: 1) the simple test is passed when the clock rate estimate for the observed data does not overlap with the range of estimates obtained from the randomized sets. 2) The intermediate test is passed when the clock rate estimate for the observed data does not overlap with the confidence intervals of the estimates obtained from the randomized sets. 3) The stringent test is passed when the confidence interval of the clock rate estimate for the observed data does not overlap with the confidence intervals of the estimates obtained from the randomized sets.

### Bayesian phylogenetic analysis

Bayesian molecular clock analyses are computationally demanding and problematic to run on large data sets. Therefore we reduced the thirteen largest data sets (MTBC, L1, L1.1.1, L1.1.1.1, L2, L2.2.1, L2.2.1.1, L2.2.1_nc1, L2.2.1_nc3, L4, L4.1.2, L4.10 and *M. bovis*) to 300 randomly selected strains. For each data set we used the Bayesian information criterion implemented in jModelTest 2.1.10 v20160303 (Darriba et al. 2012) to identify the best fitting nucleotide substitution model among 11 possible schemes including unequal nucleotide frequencies (total models = 22, options -s 11 and -f). We performed Bayesian inference with Beast2 (Bouckaert et al. 2014). We corrected the xml file to specify the number of invariant sites as indicated here: https://groups.google.com/forum/#!topic/beast-users/QfBHMOqImFE, and used the tip sampling year as calibration.

We ran four Beast analyses with different settings: we used a relaxed lognormal clock model (Drummond et al. 2006), the best fitting nucleotide substitution model according to the results of jModelTest, and two different coalescent priors: constant population size and exponential population growth (or shrinkage). We chose a 1/x prior for the population size [0- 10^9^], two different priors for the mean of the lognormal distribution of the clock rate (1/x and uniform) [10^−10^ − 10^−5^], a normal(0,1) prior for the standard deviation of the lognormal distribution of the clock rate [0 – infinity]. For the exponential growth rate prior, we used the standard Laplace distribution [−infinity − infinity]. For all data sets, we ran at least two runs, we used Tracer 1.7.1 (Rambaut et al. 2018) to identify and exclude the burn-in, to evaluate convergence among runs and to calculate the estimated sample size (ESS). We stopped the runs when at least two chains reached convergence, and the ESS of the posterior and of all parameters were larger than 200.

### Analyses with the complete MTB complex and aDNA

We analyzed the complete data set of 6,285 genomes with the same methods described above. The only difference was that for the LSD analysis, we rooted the input tree using *Mycobacterium canetti* (SAMN00102920, SRR011186) as outgroup. We did this because we noticed that without outgroup, all methods placed the root on the branch separating *M. bovis* from all other lineages, and not on the branch separating MTB *sensu stricto* from the other lineages.

To test the time dependency hypothesis, we repeated the LSD and Beast analyses on the MTB complex, adding the aDNA genome sequences of three MTB strains obtained from Precolumbian Peruvian human remains (Bos et al. 2014). These are the most ancient aDNA samples available for MTB. For LSD, we assigned as sampling year the confidence interval of the radiocarbon dating reported in the original publication. For Beast, we assigned uniform priors spanning the confidence interval but we failed to reach convergence, therefore we used the mean of the maximum and minimum years in the confidence interval (SAMN02727818: 1126 [1028-1224], SAMN02727820: 1117 [1023 – 1211], SAMN02727821: 1211 [1141 – 1280]). We ran three different analyses with Beast: we used the sub-sample of 300 strains with two different priors on the clock rate (1/x and uniform), and an independent sub-sample of 500 strain, for this last data set (500 strains) we assumed a HKY model and used a uniform prior on the clock rate (Sup. Table S3).

To summarize the results of the Beast analysis with the aDNA samples and retrieve the age of the MRCA of the individual lineages, we considered the analysis performed on the subset of 500 strains: we randomly sampled 5,000 trees from the posterior (after excluding the burn-in), and calculated the Maximum clade credibility tree with the software Treeannotator v2.5.0.

## Acknowledgments

Calculations were performed at sciCORE (http://scicore.unibas.ch/) scientific computing core facility at the University of Basel. This work was supported by the Swiss National Science Foundation (grants 310030_166687, IZRJZ3_164171, IZLSZ3_170834 and CRSII5_177163), the European Research Council (309540-EVODRTB) and SystemsX.ch. SD was supported by a McKenzie fellowship from the University of Melbourne.

## Supplementary Tables and Figures

**Supplementary Table S1**

File: Supplementary_tableS1.tsv

List of strains used in this study with sampling year and accession numbers

**Supplementary Figure S2.**
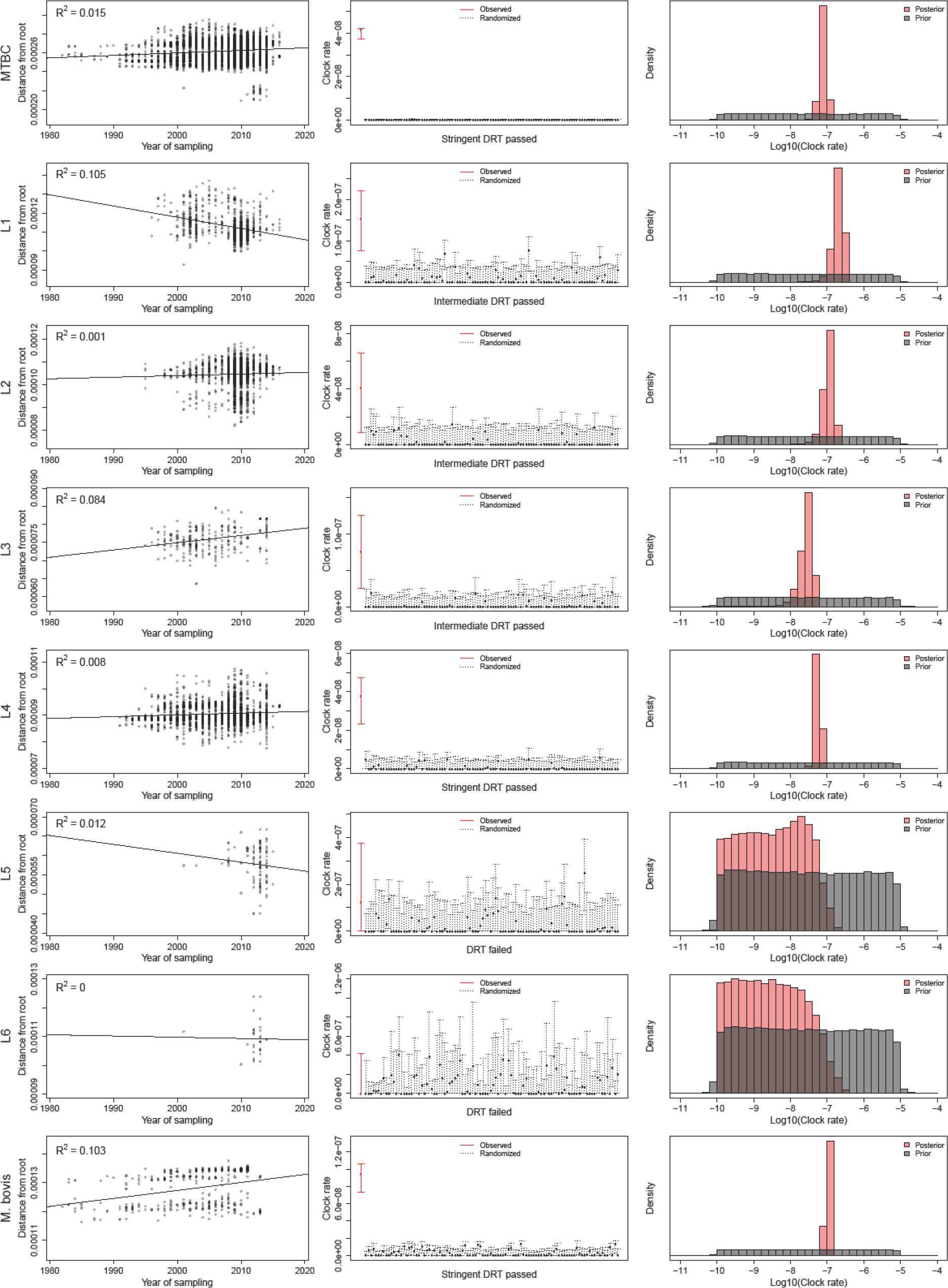
For each data set we report the results of the root to tip regression, the results of the Date Randomization Test (DRT) with LSD, and the comparison of the prior and posterior distribution of the clock rate. The simple DRT is passed when the clock rate estimate for the observed data does not overlap with the range of estimates obtained from the randomized sets. The intermediate DRT is passed when the clock rate estimate for the observed data does not overlap with the confidence intervals of the estimates obtained from the randomized sets. The stringent DRT is passed when the confidence interval of the clock rate estimate for the observed data does not overlap with the confidence intervals of the estimates obtained from the randomized sets. Large data sets (MTBC, L1, L2, L4 and *M. bovis*) were randomly sub-sampled to 300 strains for the Beast analysis.

**Supplementary Table S3**

File: Supplementary_tableS3.xlsx

Results of Beast and LSD for all data sets

**Supplementary Figure S4.**
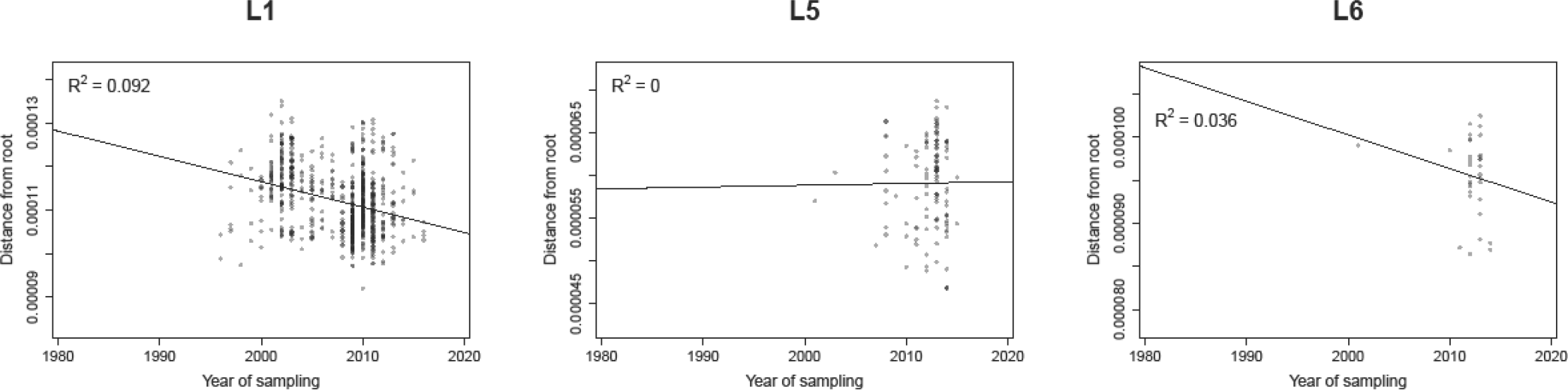
Root to tip regression analysis of L1, L5 and L6. The difference compared to Supplementary Fig. S2 is that the root was not placed in the position that minimizes the sum of the squared residuals from the regression line, but was obtained from the complete MTBC tree as shown in Fig. 3a, and it is therefore defined by the outgroup of each of these lineages.

**Supplementary Figure S5.**
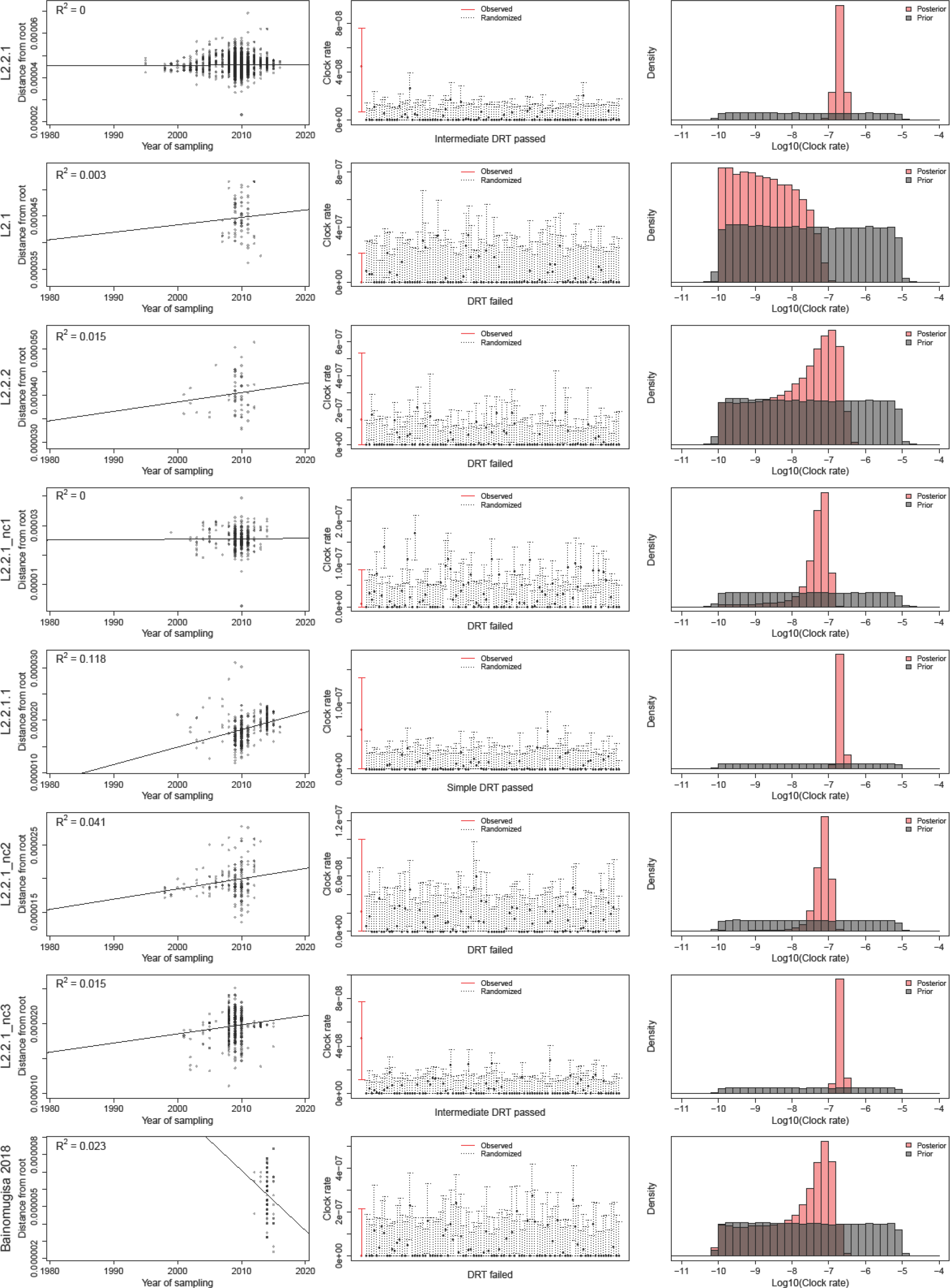
For each data set we report the results of the root to tip regression, the results of the Date Randomization Test (DRT) with LSD, and the comparison of the prior and posterior distribution of the clock rate. The simple DRT is passed when the clock rate estimate for the observed data does not overlap with the range of estimates obtained from the randomized sets. The intermediate DRT is passed when the clock rate estimate for the observed data does not overlap with the confidence intervals of the estimates obtained from the randomized sets. The stringent DRT is passed when the confidence interval of the clock rate estimate for the observed data does not overlap with the confidence intervals of the estimates obtained from the randomized sets. Large data sets (L1.1.1, L1.1.1.1, L2.2.1, L2.2.1.1, L2.2.1_nc1, L2.2.1_nc3, L4.10, L4.1.2) were randomly sub-sampled to 300 strains for the Beast analysis.

**Supplementary Figure S6.**
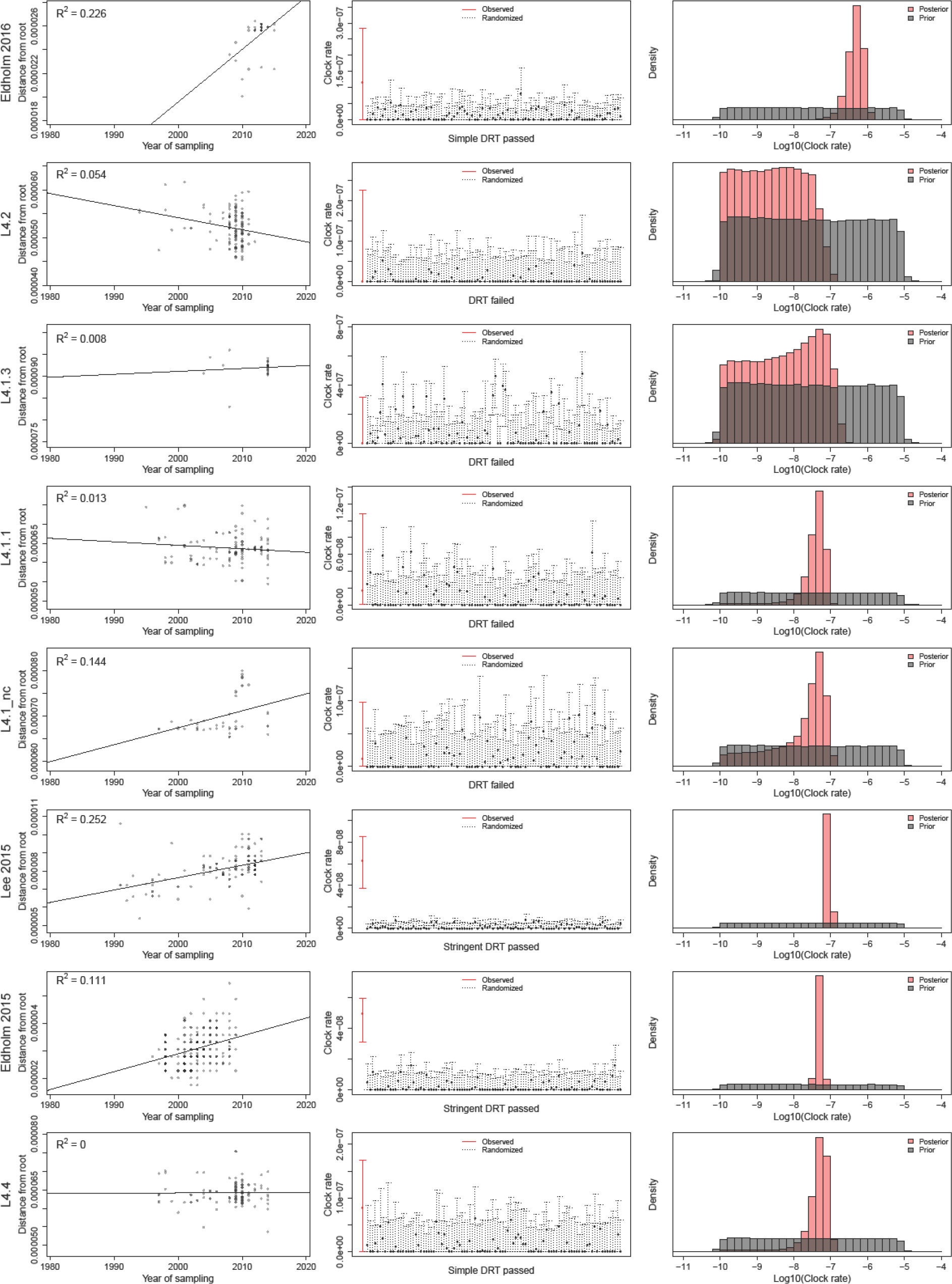
For each data set we report the results of the root to tip regression, the results of the Date Randomization Test (DRT) with LSD, and the comparison of the prior and posterior distribution of the clock rate. The simple DRT is passed when the clock rate estimate for the observed data does not overlap with the range of estimates obtained from the randomized sets. The intermediate DRT is passed when the clock rate estimate for the observed data does not overlap with the confidence intervals of the estimates obtained from the randomized sets. The stringent DRT is passed when the confidence interval of the clock rate estimate for the observed data does not overlap with the confidence intervals of the estimates obtained from the randomized sets. Large data sets (L1.1.1, L1.1.1.1, L2.2.1, L2.2.1.1, L2.2.1_nc1, L2.2.1_nc3, L4.10, L4.1.2) were randomly sub-sampled to 300 strains for the Beast analysis.

**Supplementary Figure S7.**
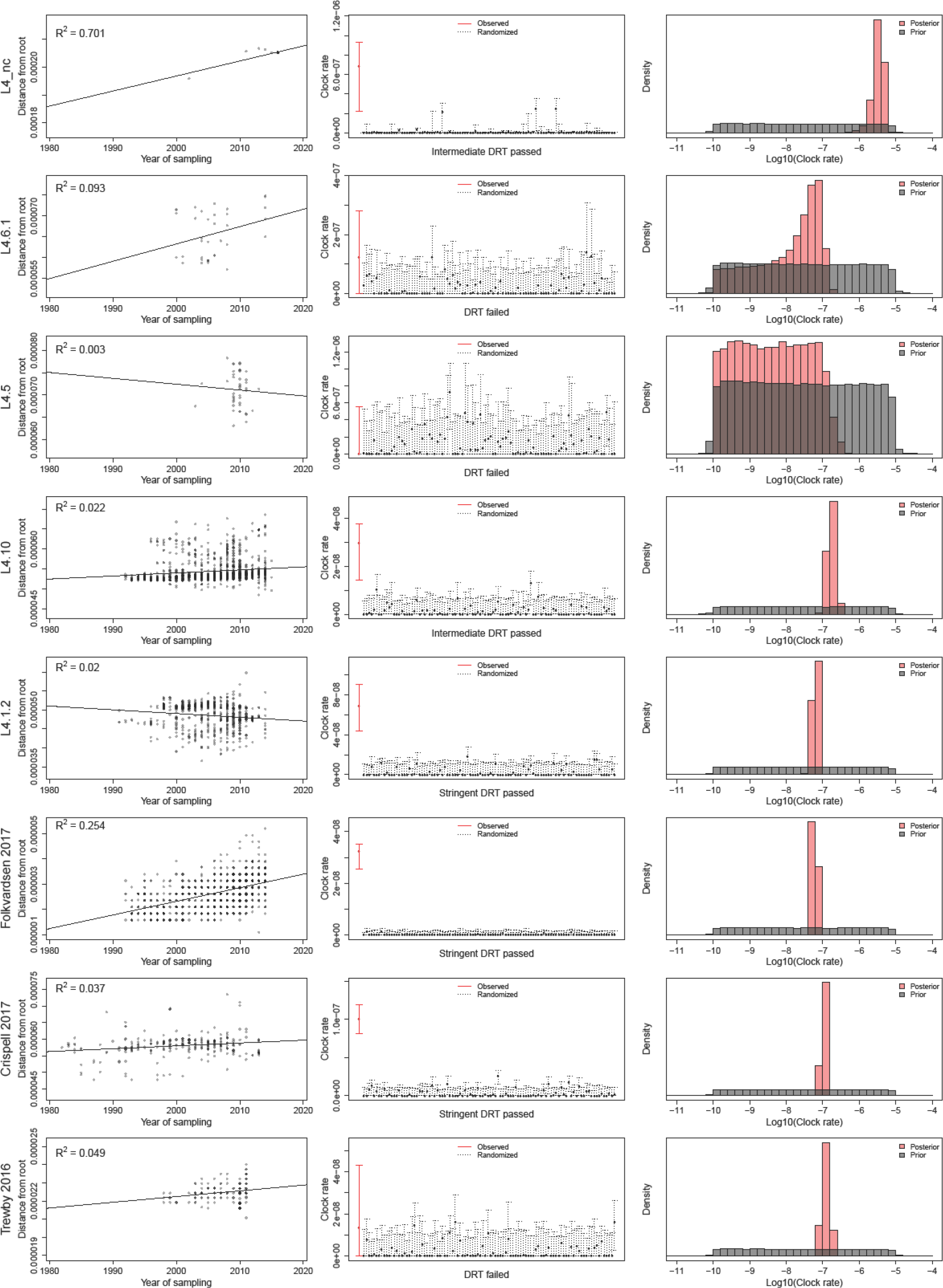
For each data set we report the results of the root to tip regression, the results of the Date Randomization Test (DRT) with LSD, and the comparison of the prior and posterior distribution of the clock rate. The simple DRT is passed when the clock rate estimate for the observed data does not overlap with the range of estimates obtained from the randomized sets. The intermediate DRT is passed when the clock rate estimate for the observed data does not overlap with the confidence intervals of the estimates obtained from the randomized sets. The stringent DRT is passed when the confidence interval of the clock rate estimate for the observed data does not overlap with the confidence intervals of the estimates obtained from the randomized sets. Large data sets (L1.1.1, L1.1.1.1, L2.2.1, L2.2.1.1, L2.2.1_nc1, L2.2.1_nc3, L4.10, L4.1.2) were randomly sub-sampled to 300 strains for the Beast analysis.

**Supplementary Figure S8.**
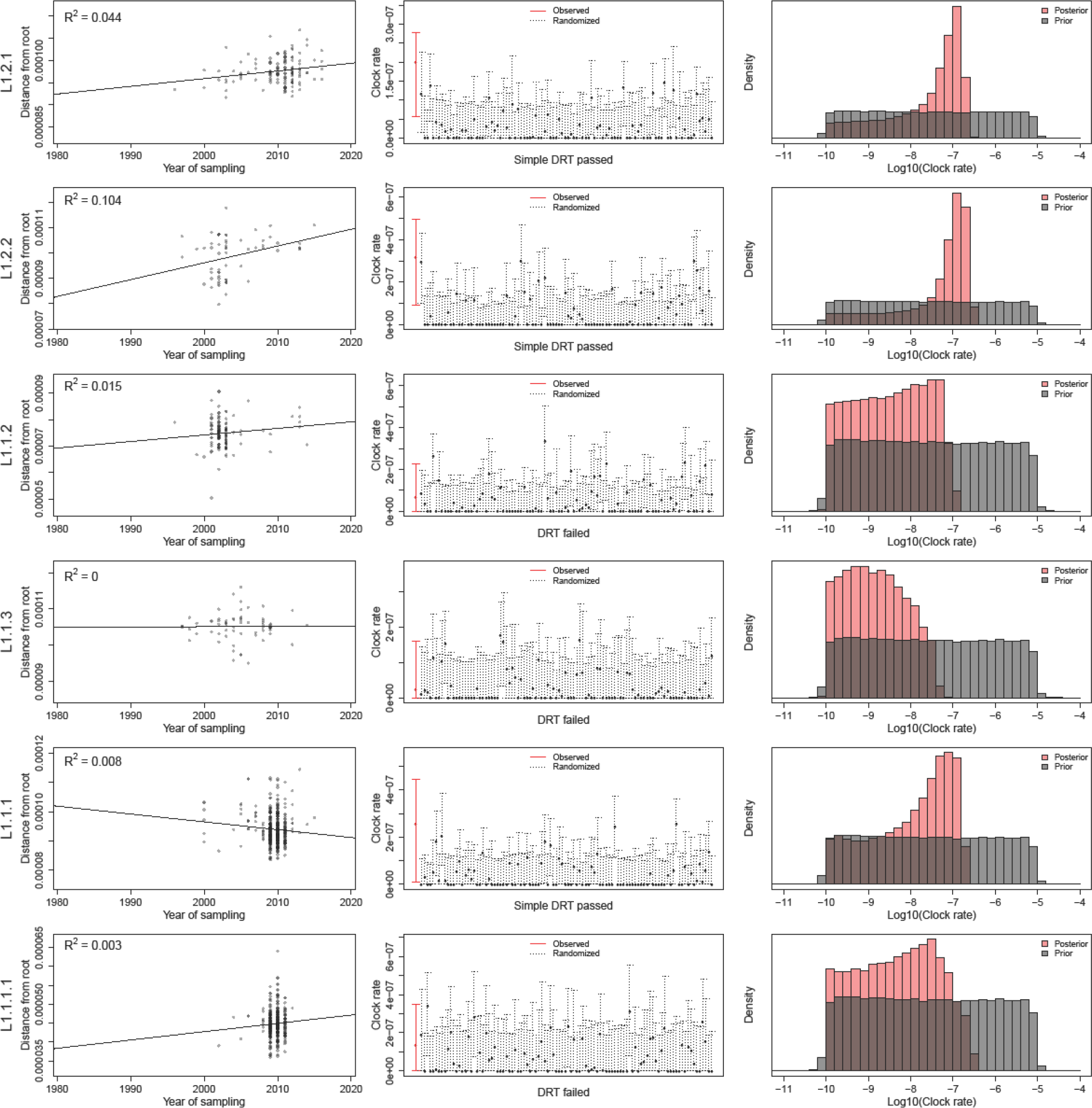
For each data set we report the results of the root to tip regression, the results of the Date Randomization Test (DRT) with LSD, and the comparison of the prior and posterior distribution of the clock rate. The simple DRT is passed when the clock rate estimate for the observed data does not overlap with the range of estimates obtained from the randomized sets. The intermediate DRT is passed when the clock rate estimate for the observed data does not overlap with the confidence intervals of the estimates obtained from the randomized sets. The stringent DRT is passed when the confidence interval of the clock rate estimate for the observed data does not overlap with the confidence intervals of the estimates obtained from the randomized sets. Large data sets (L1.1.1, L1.1.1.1, L2.2.1, L2.2.1.1, L2.2.1_nc1, L2.2.1_nc3, L4.10, L4.1.2) were randomly sub-sampled to 300 strains for the Beast analysis.

**Supplementary Figure S9.**
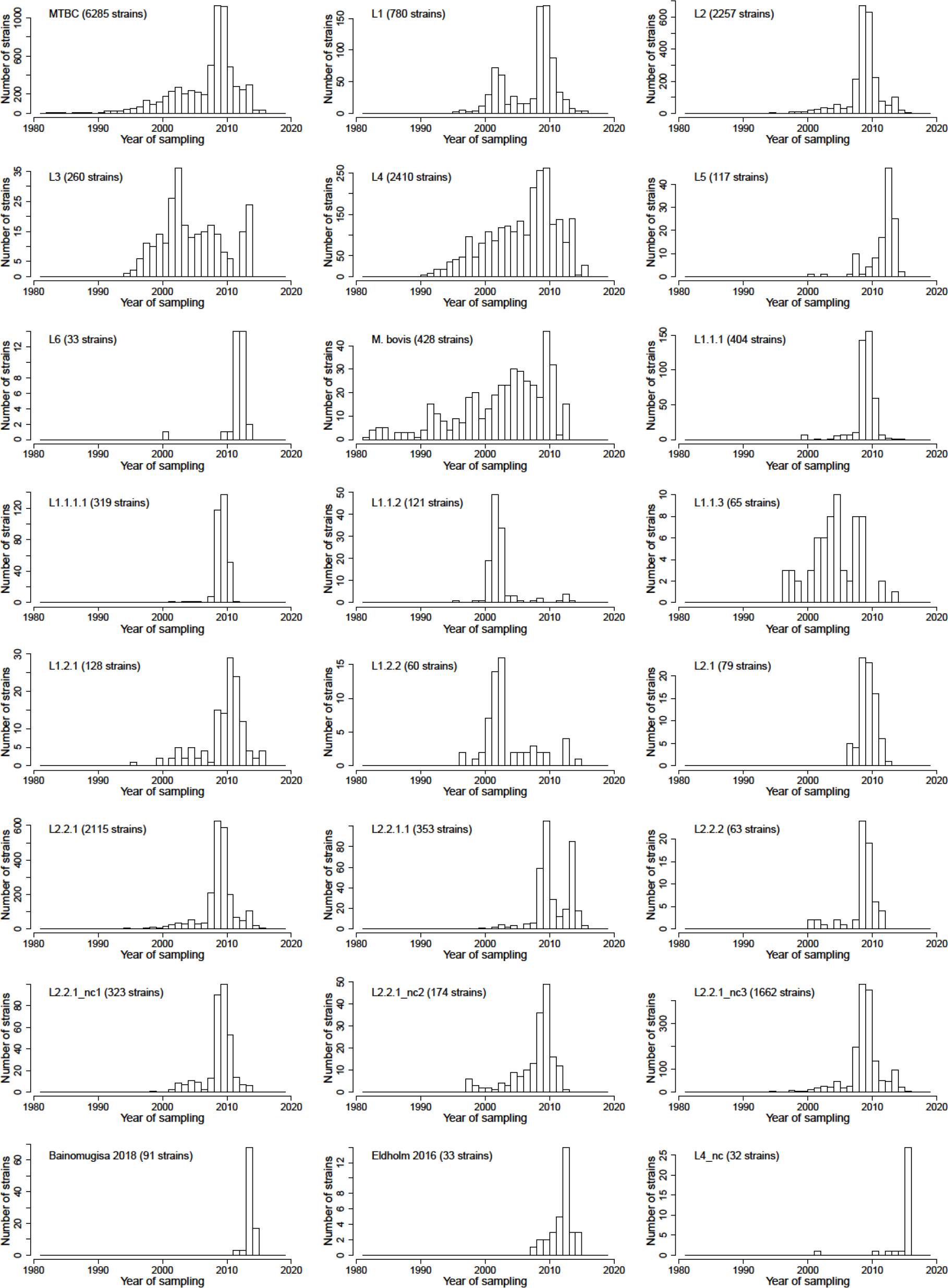
Distribution of the sampling years for all data sets

**Supplementary Figure S10.**
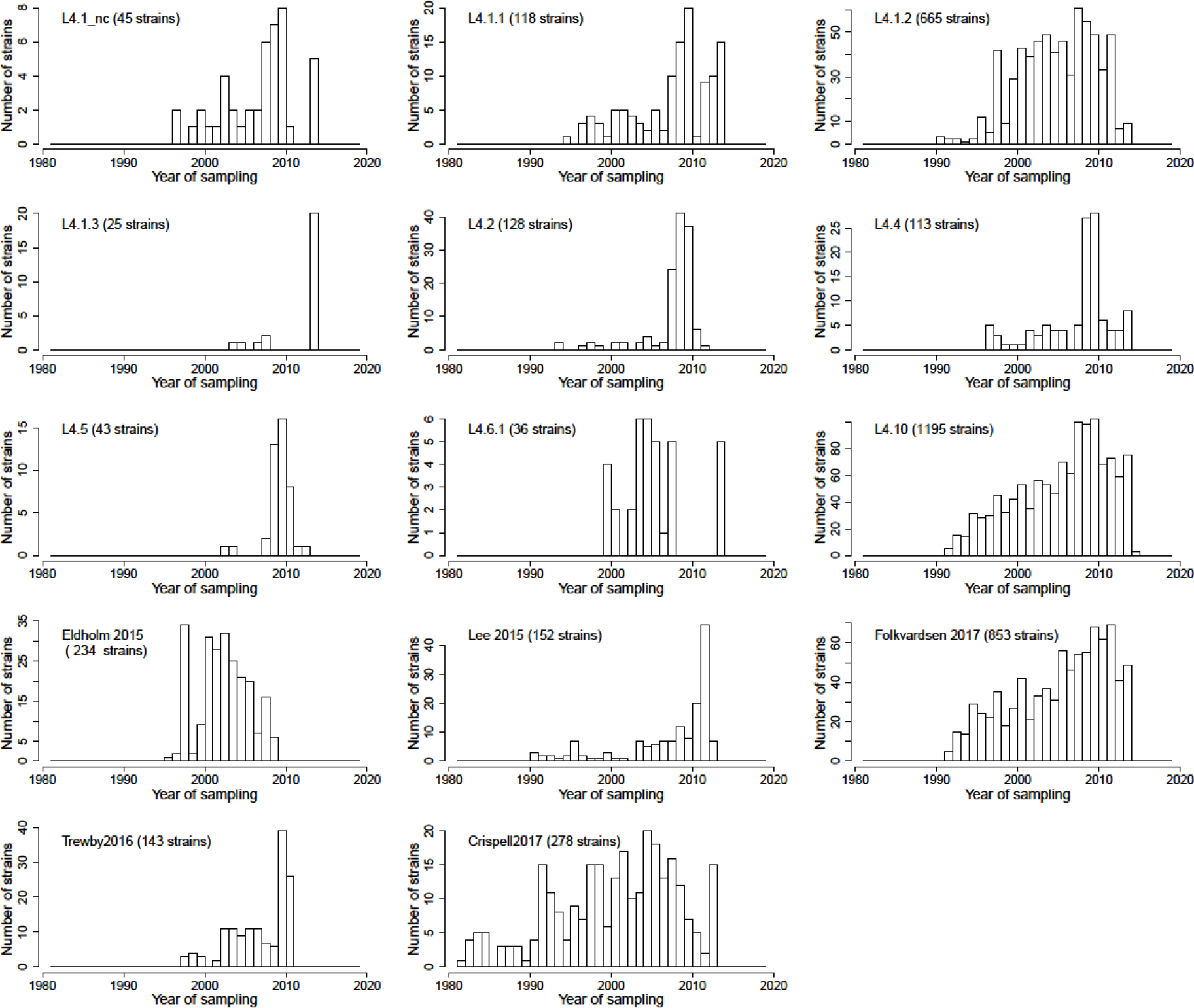
Distribution of the sampling years for all data sets

**Supplementary Figure S11.**
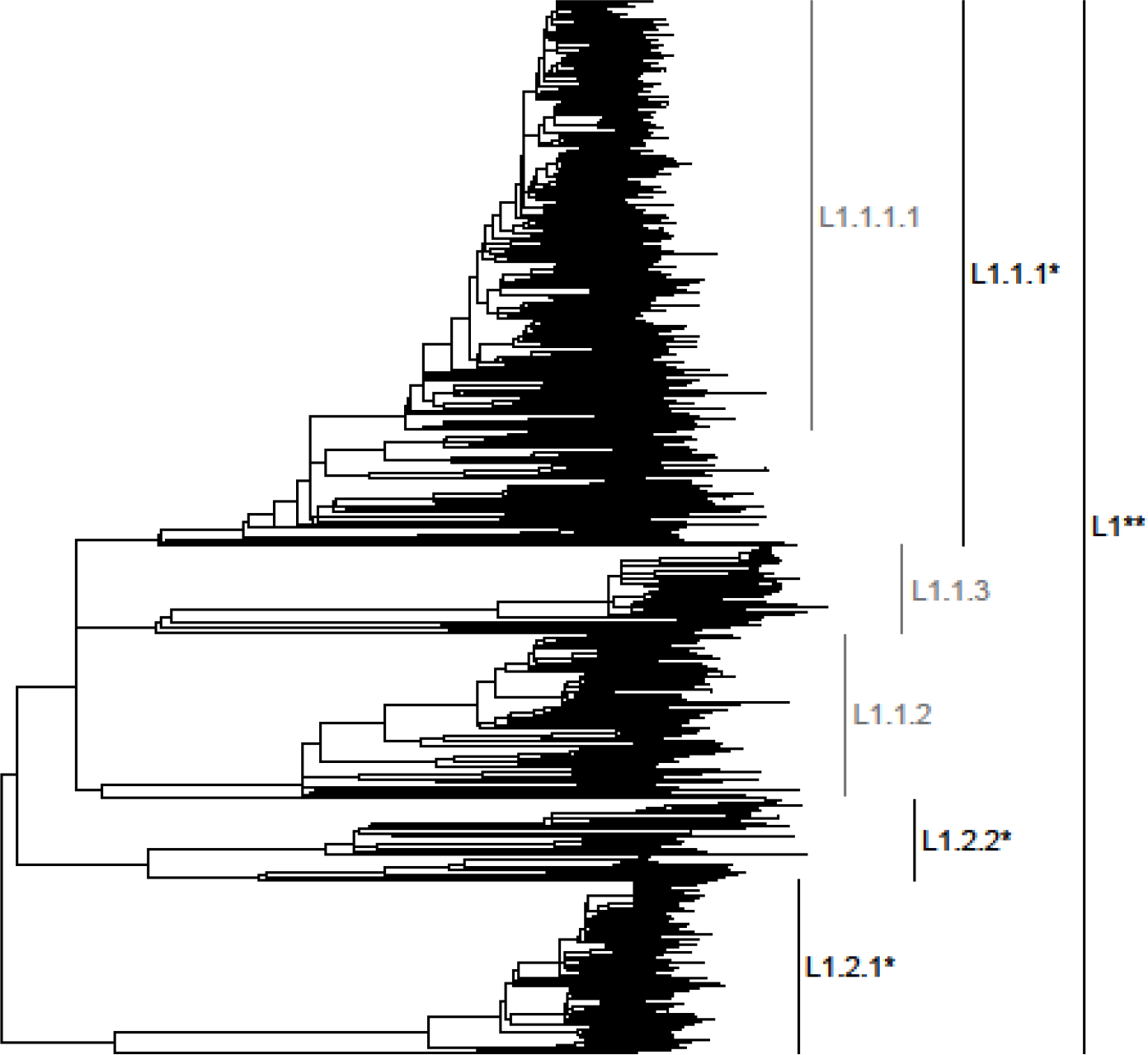
Sub-lineages of L1 that were included in the analysis. Clades colored in gray did not pass the DRT, clades colored in black passed the DRT. *: simple DRT passed, ** intermediate DRT passed, ***: stringent DRT passed.

**Supplementary Figure S12.**
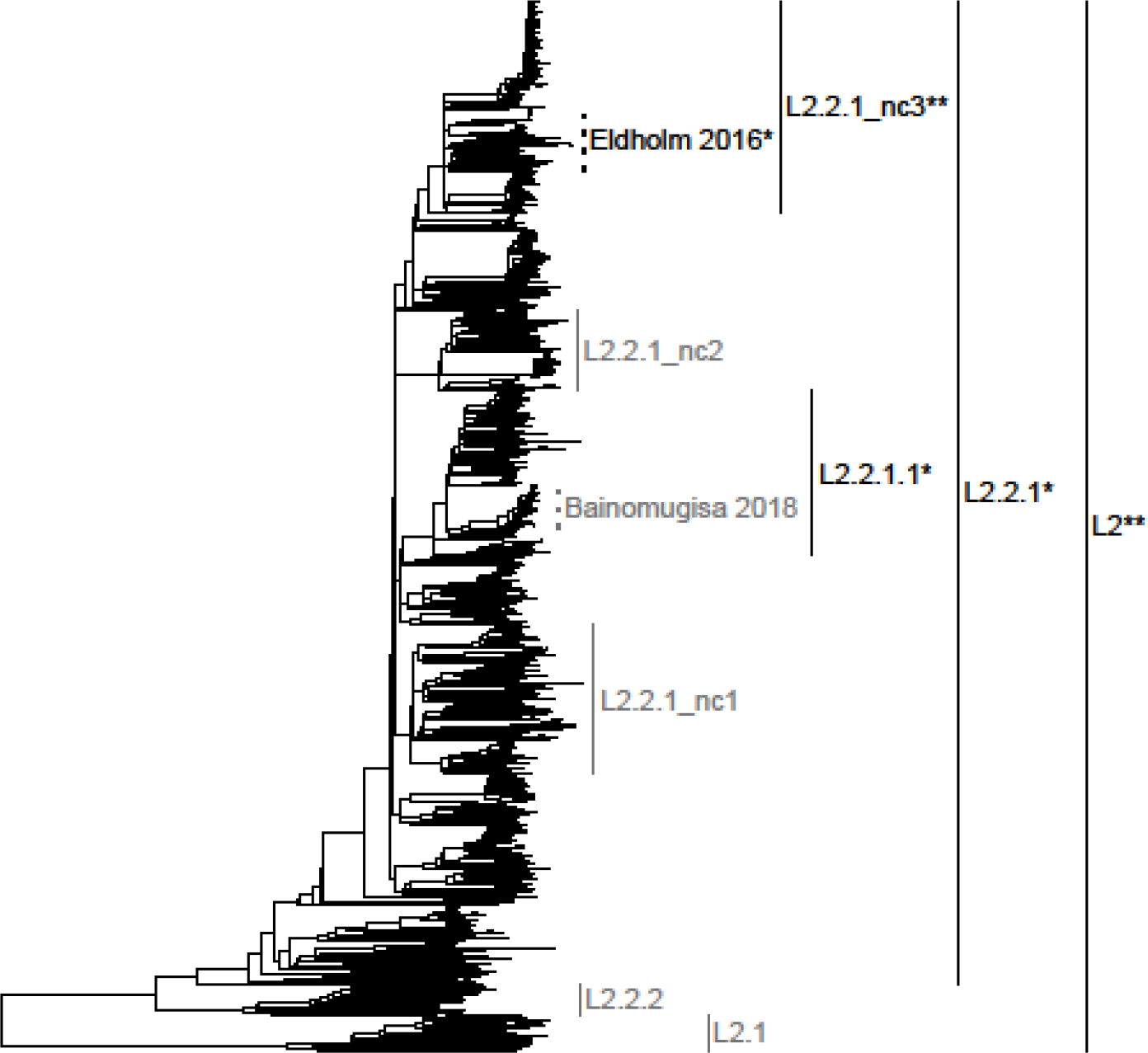
Sub-lineages and outbreaks of L2 that were included in the analysis. Clades colored in gray did not pass the DRT, clades colored in black passed the DRT. *: simple DRT passed, ** intermediate DRT passed, ***: stringent DRT passed. Dotted lines represent local two outbreaks from previous studies.

**Supplementary Figure S13.**
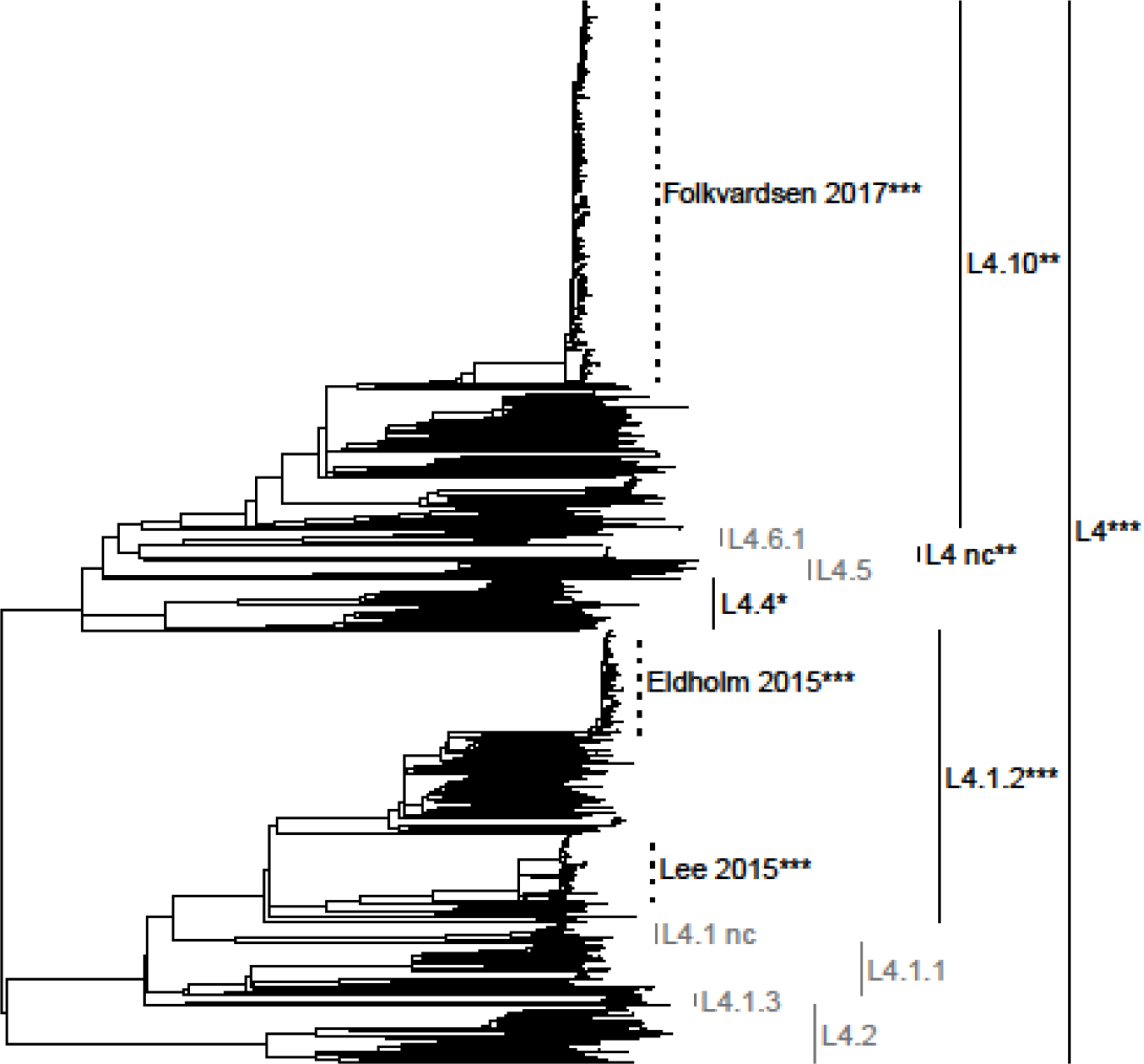
Sub-lineages and outbreaks of L4 that were included in the analysis. Clades colored in gray did not pass the DRT, clades colored in black passed the DRT. *: simple DRT passed, ** intermediate DRT passed, ***: stringent DRT passed. Dotted lines represent three outbreaks from previous studies.

**Supplementary Figure S14.**
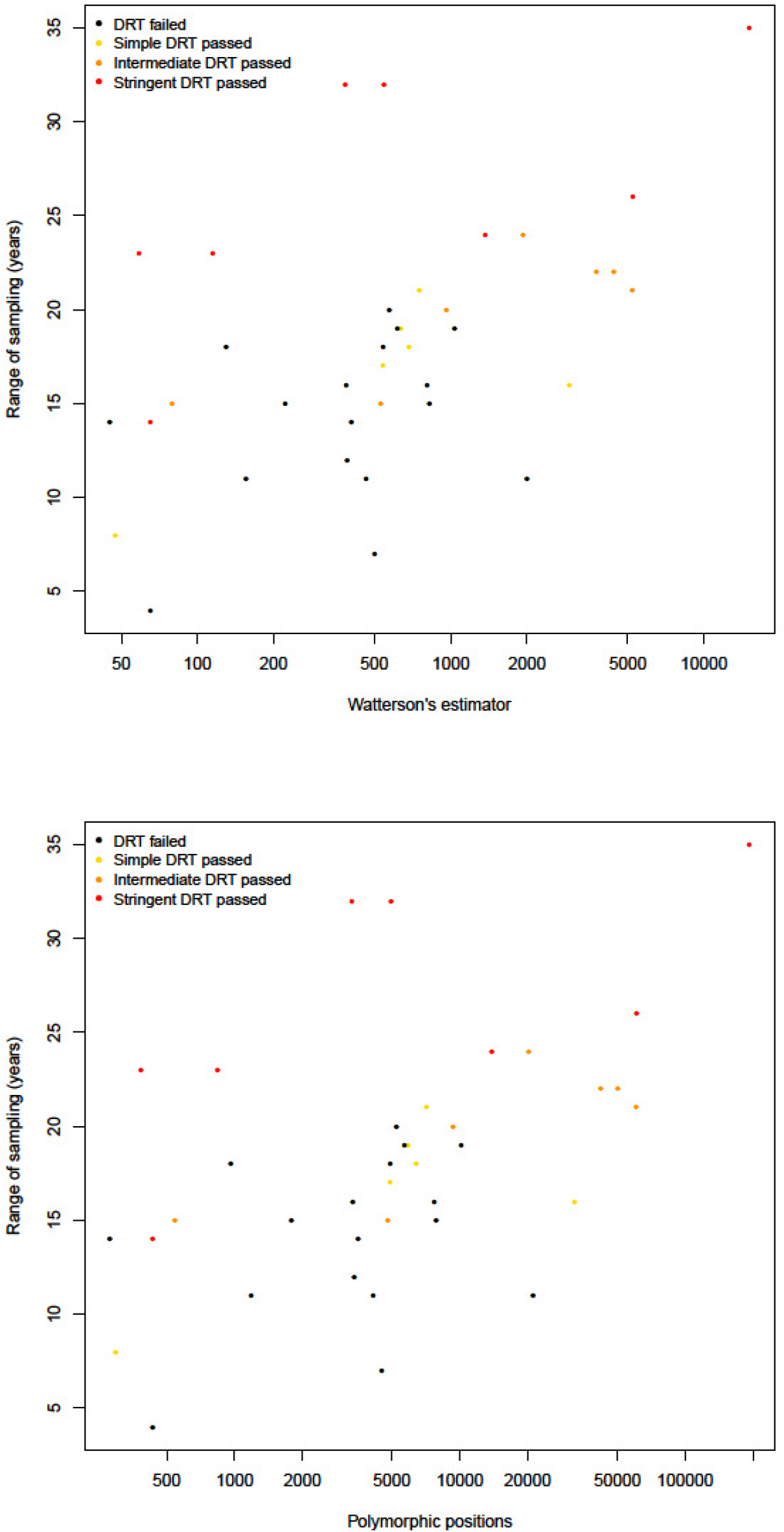
Results of the DRT for all data sets ordered by genetic diversity (Watterson’s estimator and number of polymorphic positions) and temporal range. Data sets with fewer strains sampled in a shorter period of time tended to fail the DRT irrespectively of the genetic diversity of the data set.

**Supplementary Figure S15.**
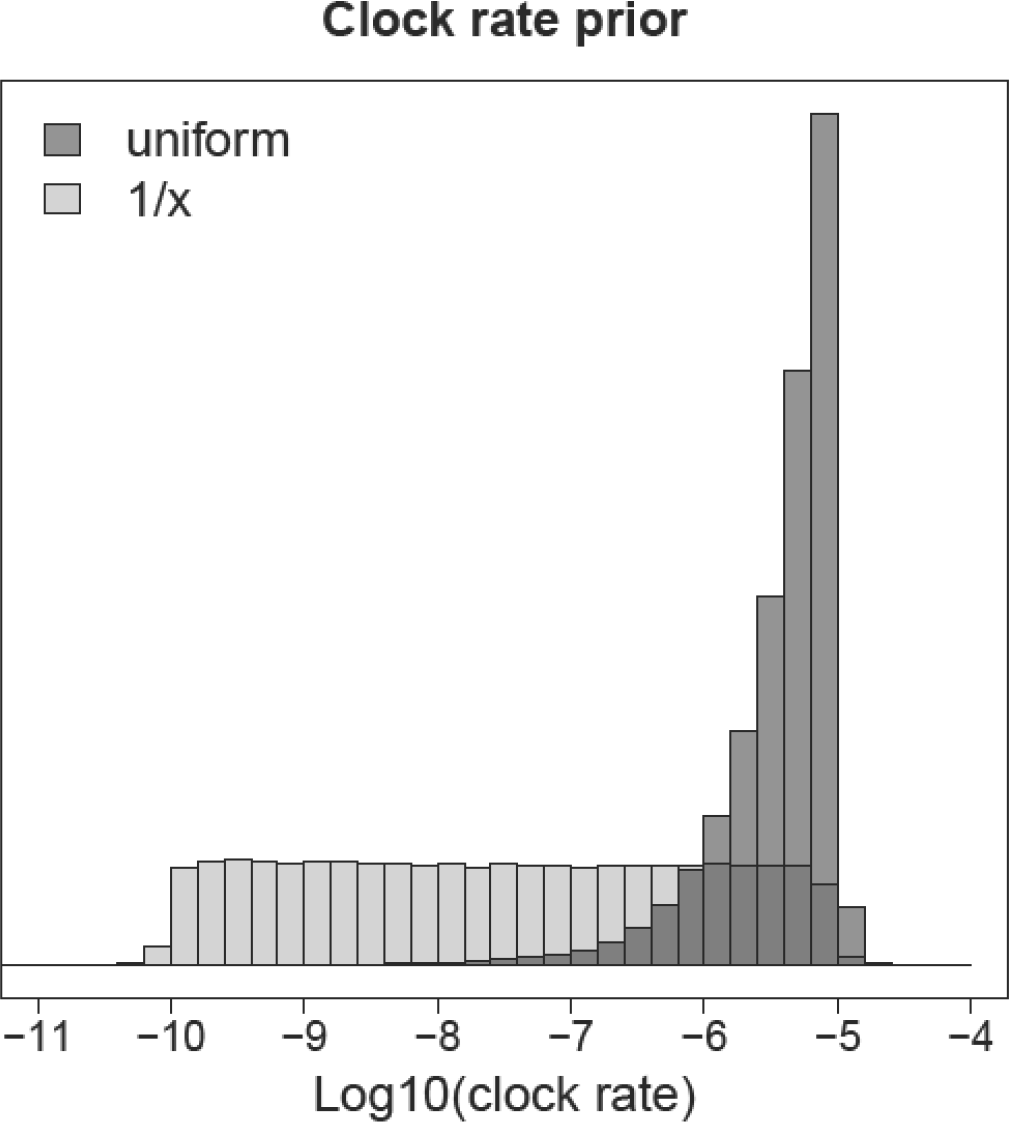
Comparison of different priors on the clock rate (1/x prior and uniform prior). The uniform prior place most weight on high clock rates, while the 1/x prior distributes the weight through all order of magnitude.

**Supplementary Figure S16.**
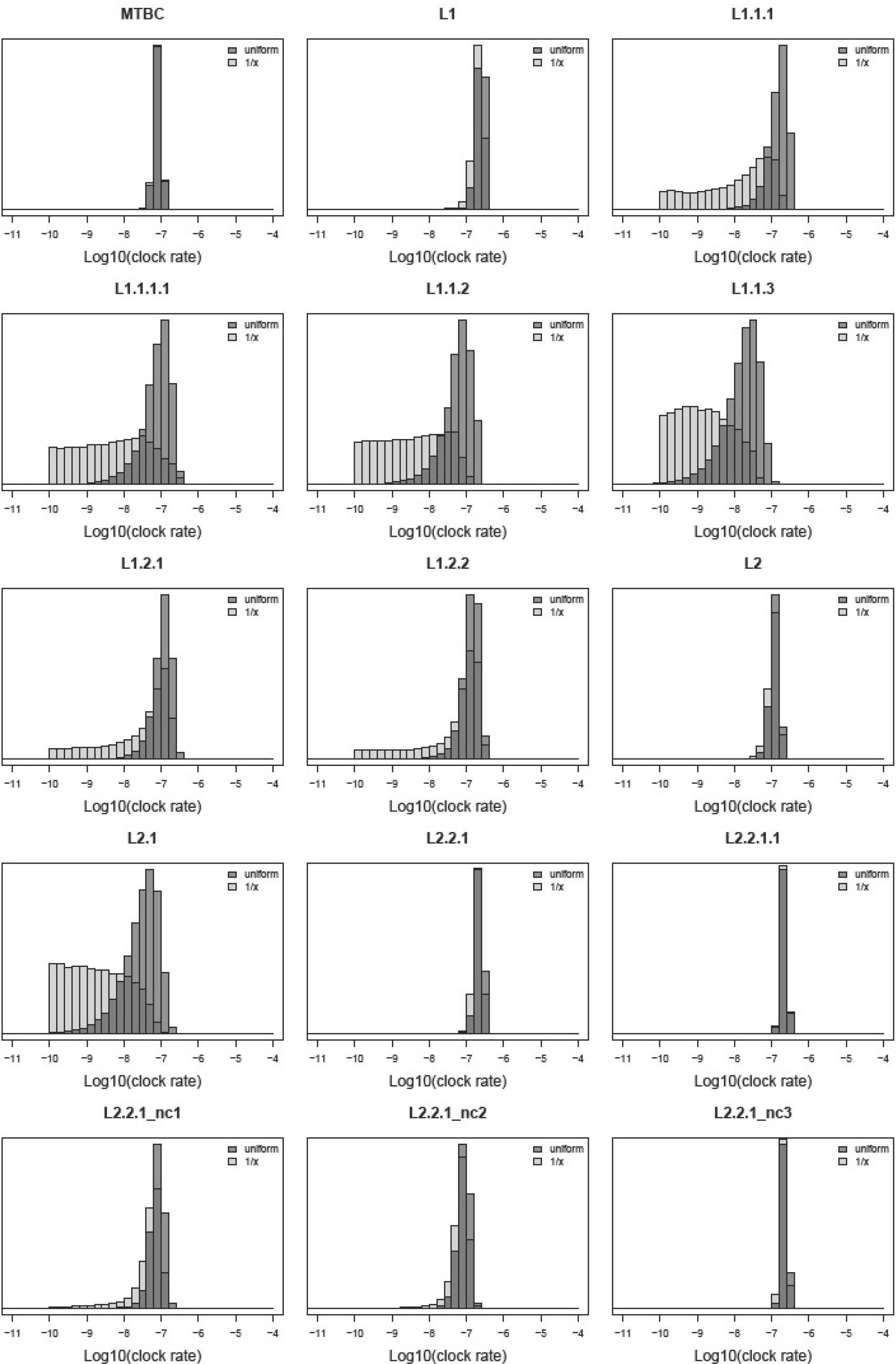
Posterior distribution of the clock rate, obtained with two different priors (1/x and uniform [10^−10^ – 10^−5^]). The prior distributions for the two analyses are shown in Sup. Fig. S15.

**Supplementary Figure S17.**
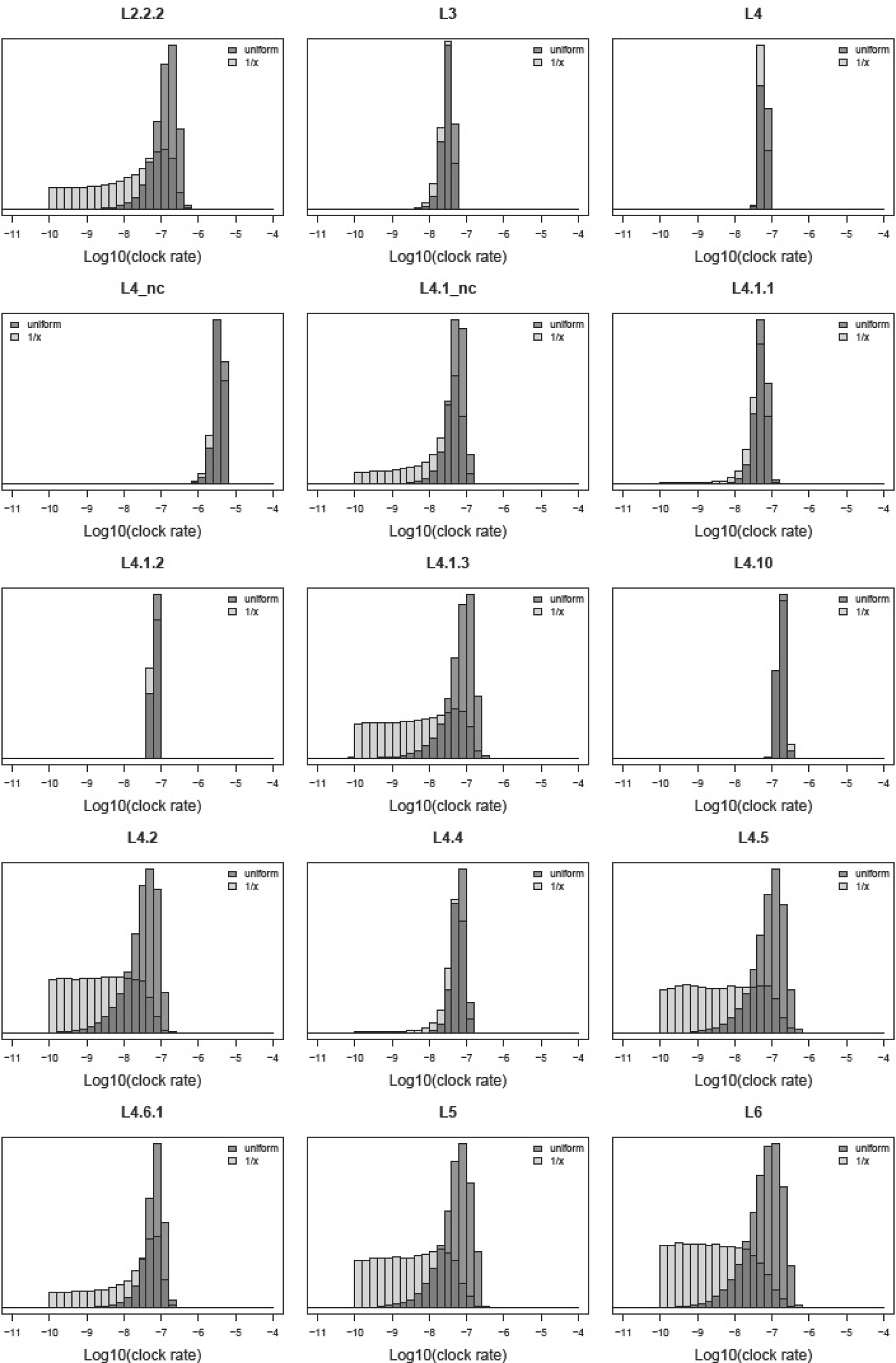
Posterior distribution of the clock rate, obtained with two different priors (1/x and uniform [10^−10^ – 10^−5^]). The prior distributions for the two analyses are shown in Sup. Fig. S15.

**Supplementary Figure S18.**
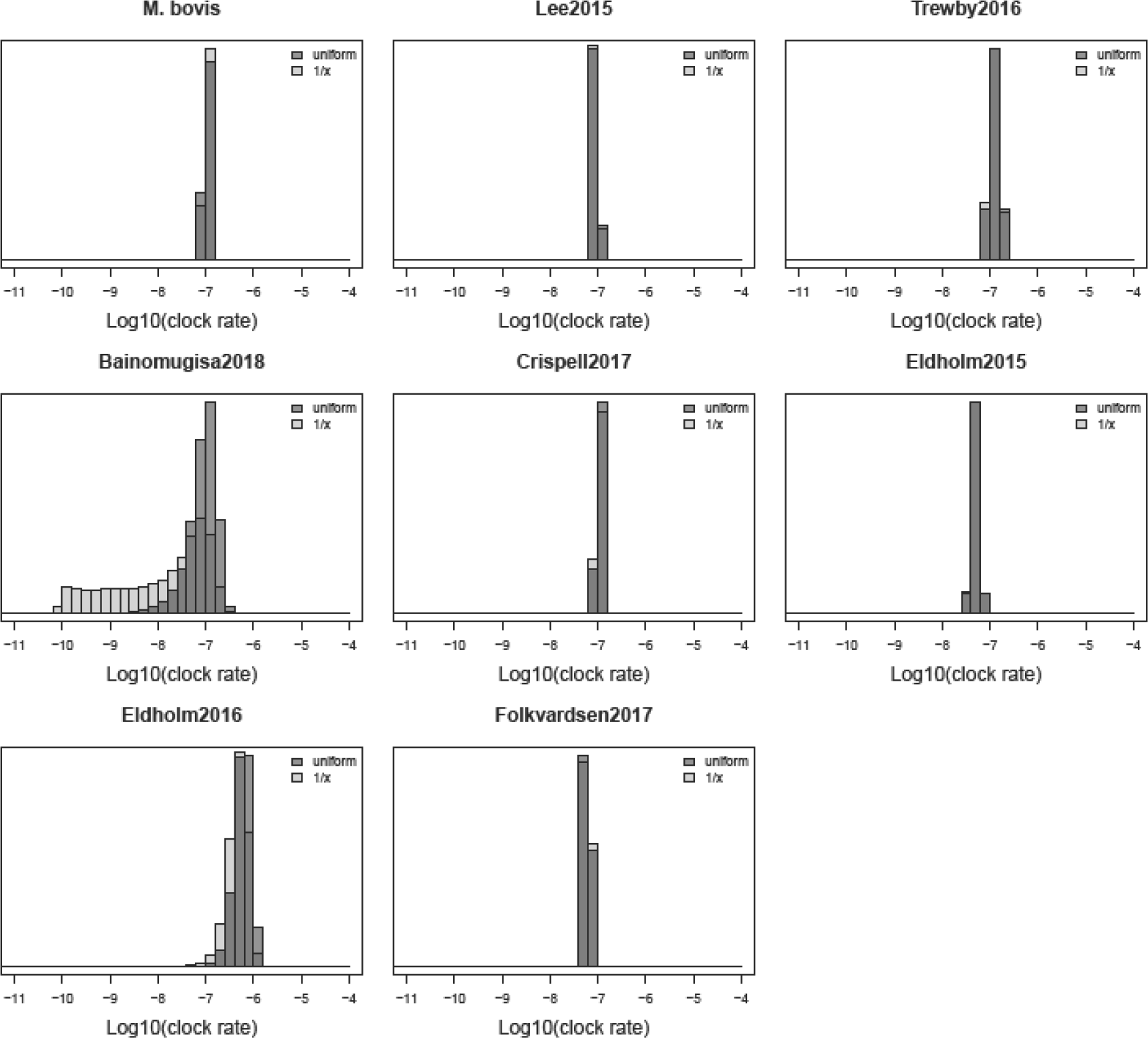
Posterior distribution of the clock rate, obtained with two different priors (1/x and uniform [10^−10^ – 10^−5^]). The prior distributions for the two analyses are shown in Sup. Fig. S15.

**Supplementary Figure S19.**
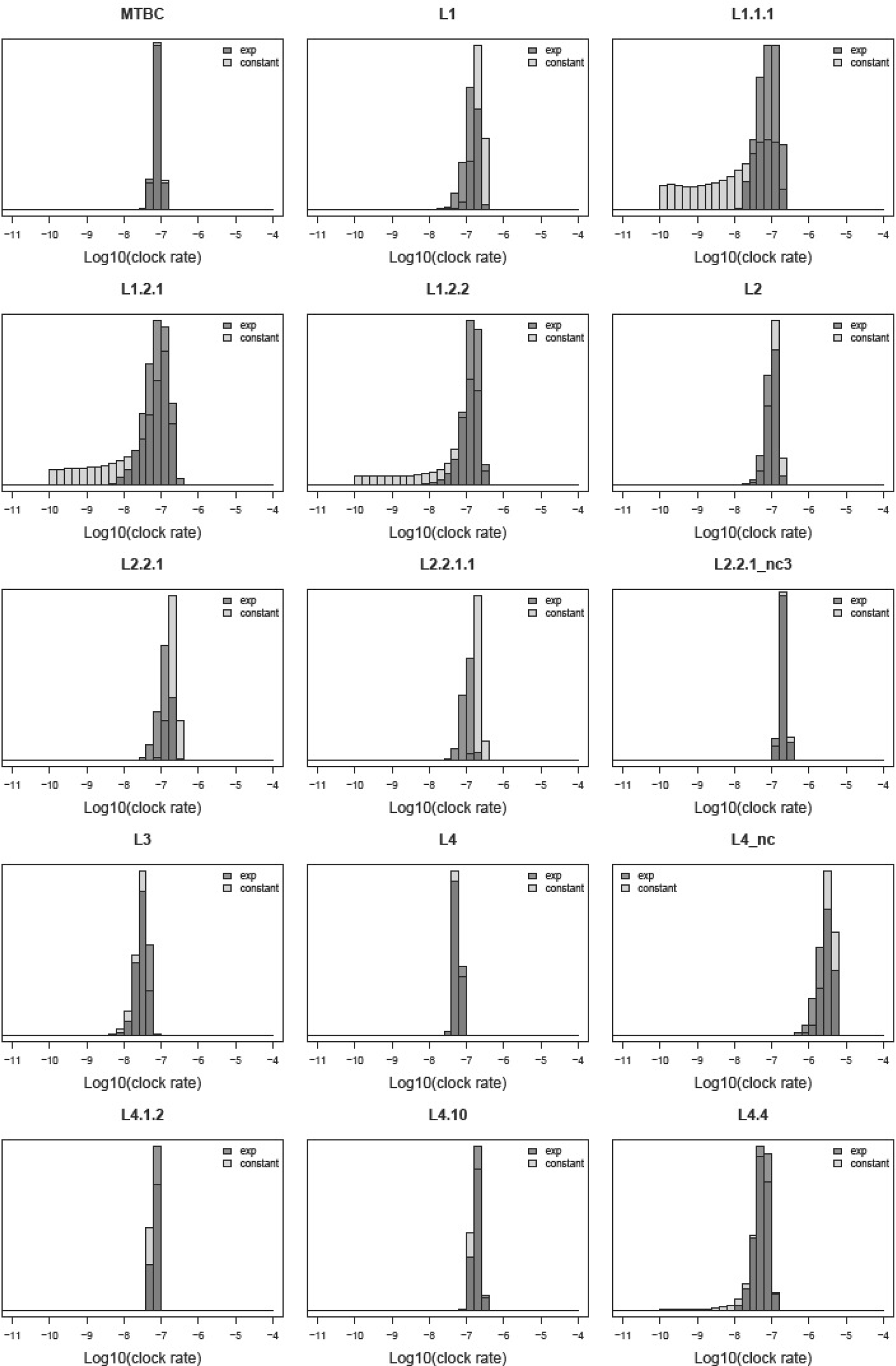
Comparison of the posterior distribution of the clock rate obtained with a constant population size and an exponential population growth prior.

**Supplementary Figure S20.**
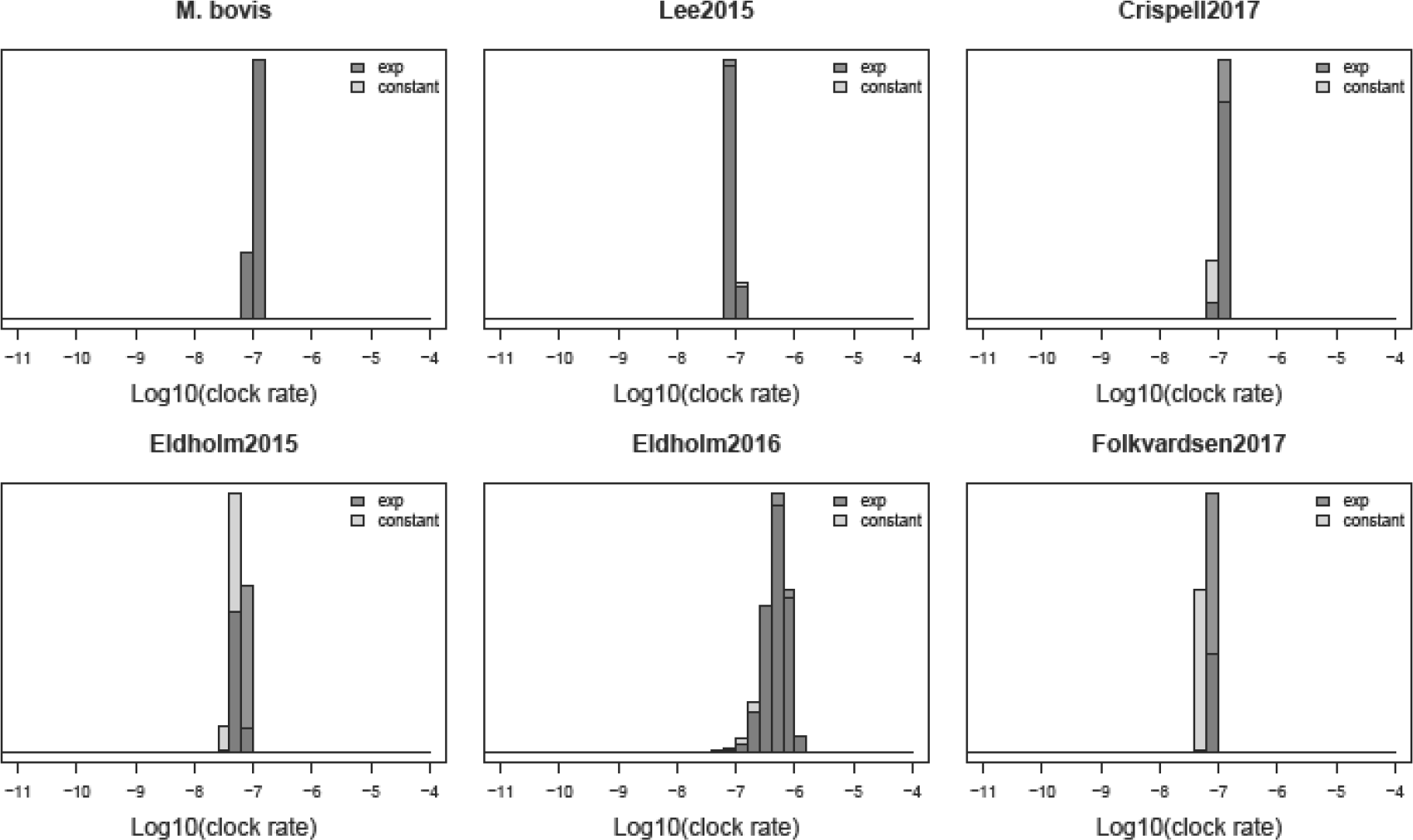
Comparison of the posterior distribution of the clock rate obtained with a constant population size and an exponential population growth prior.

**Supplementary Figure S21.**
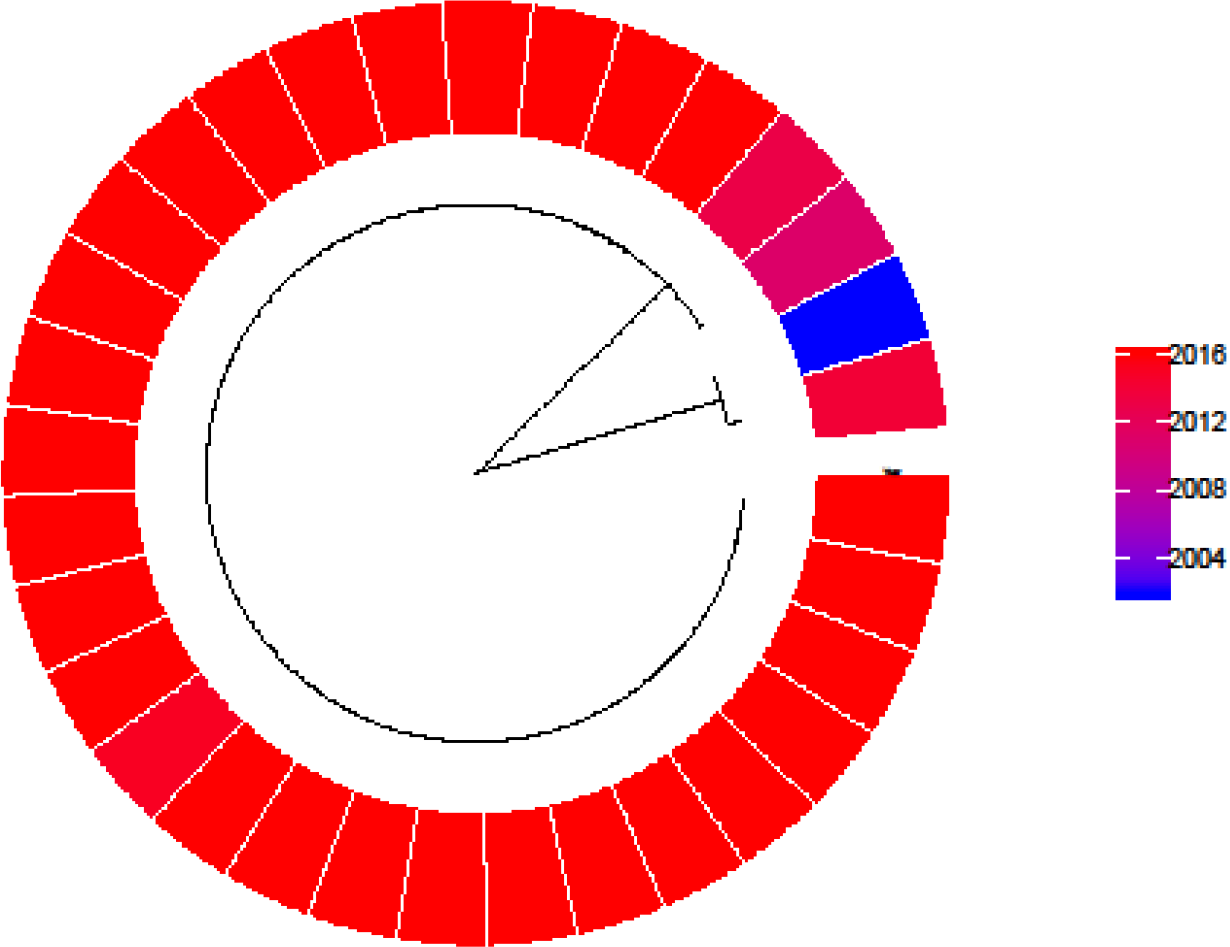
Phylogenetic tree of data set L4_nc with tips colored according to the year of sampling

**Supplementary Figure S22.**
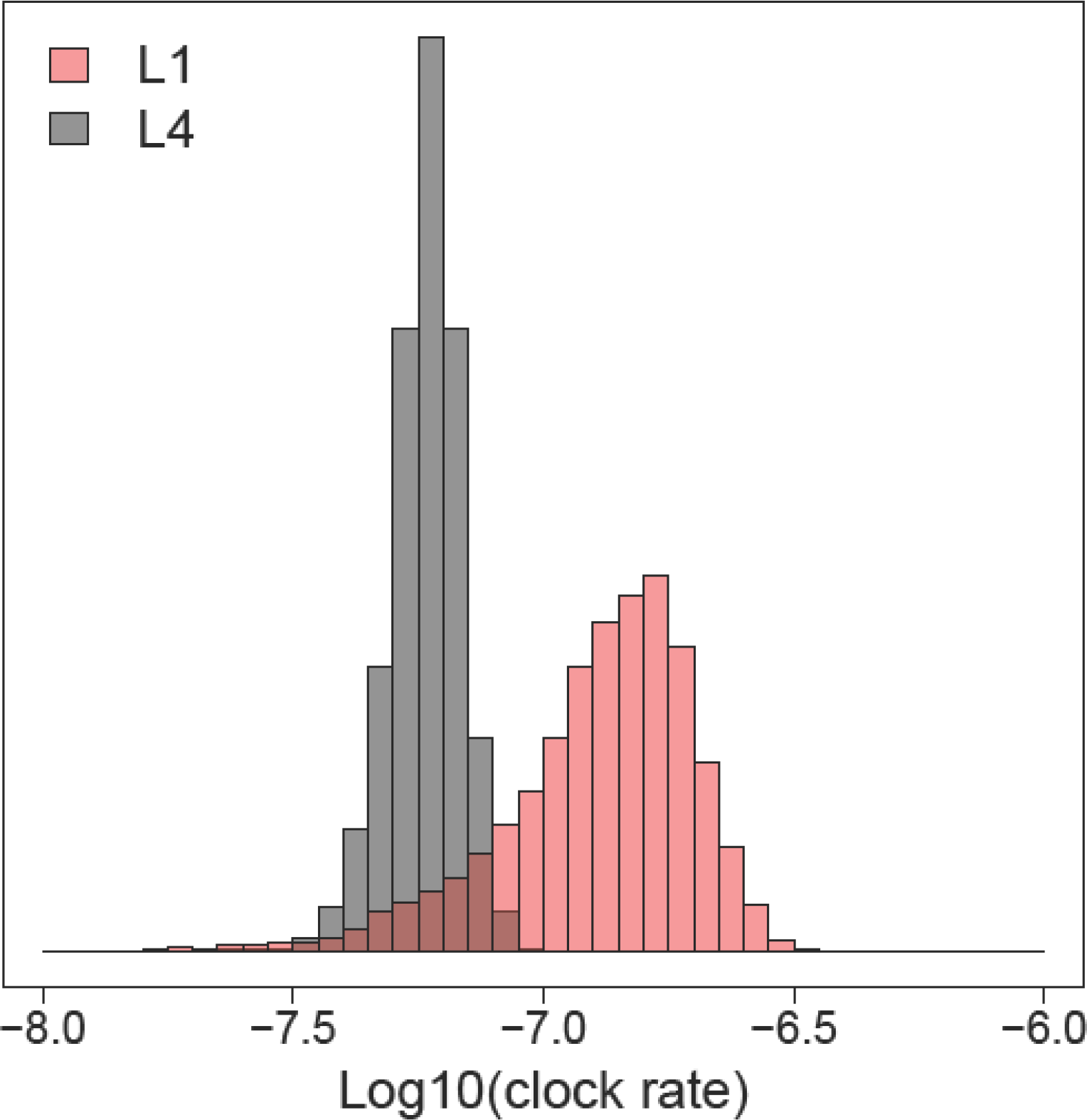
Posterior distribution of the clock rate for L1 and L4. These are the results of the analysis with the 1/x prior on the clock rate and the exponential population growth (or shrinkage) prior.

**Supplementary Table S23.**
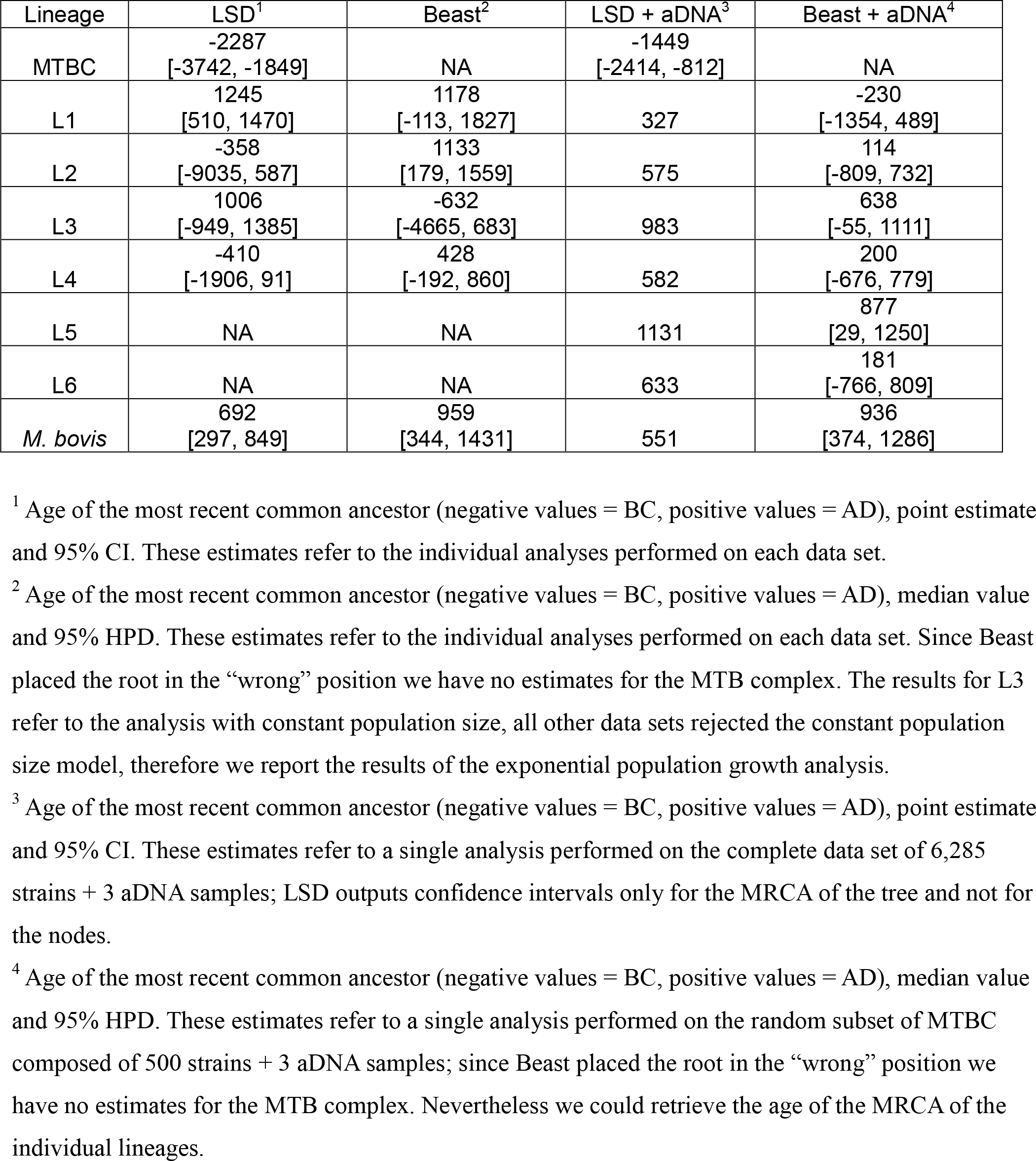
The age of the MTB complex and of its lineages resulted from different analyses

**Supplementary Table S24**

File: Supplementary_tableS24.tsv

List of all accession numbers, before filtering

